# VapA/Scs2 sustains polarized growth in *Aspergillus nidulans* by maintaining AP-2-mediated apical endocytosis

**DOI:** 10.1101/2025.07.16.665121

**Authors:** Xenia Georgiou, Sofia Politi, Sotiris Amillis, George Diallinas

## Abstract

Growth of filamentous fungi is highly polarized requiring the coordinated apical delivery of cell wall components and plasma membrane (PM) material, primarily lipids and proteins, to hyphal tips via conventional vesicular secretion. Fungal growth also requires the tight coordination of exocytosis (secretion) with endocytosis and recycling of proteins and lipids, which occurs in a defined region behind the growing tip known as the endocytic collar. Here, we genetically characterized proteins tentatively implicated in the formation of endoplasmic reticulum–plasma membrane (ER–PM) contact sites, including Scs2/VAP, tricalbins and Ist2 homologues, in *Aspergillus nidulans*. We showed that among these proteins, only the single Scs2/VapA homologue is essential for normal fungal growth, and this requirement is due to the critical role of VapA in maintaining the polarized localization of apical cargoes, such as the lipid flippases DnfA and DnfB or the SNARE protein SynA. In Δ*vapA* mutants, these cargoes lose their polarized localization, a phenotype that correlates with the mislocalization of the AP-2 cargo adaptor complex, which is essential for the endocytosis and recycling of apical membrane components. Further analysis provides evidence linking the defect in apical cargo endocytosis observed in Δ*vapA* mutants to altered membrane lipid partitioning, suggesting that VapA contributes to lipid domain organization critical for cargo recycling. Strikingly, deletion of VapA does not impair the localization or endocytosis of non-polarized (subapical) plasma membrane transporters, indicating that the trafficking and biogenesis of polarized (apical) versus non-polarized (subapical) cargoes are differentially dependent on membrane lipid composition and domain-specific organization.

## INTRODUCTION

In eukaryotic cells, the endoplasmic reticulum (ER) forms a dynamic membrane network that establishes close contacts with nearly all subcellular organelles and the plasma membrane (PM). These specialized regions, known as membrane contact sites (MCS), play a critical role in the non-vesicular transport of membrane lipids from the ER—the primary site of lipid synthesis—to other cellular compartments [1–5]. Lipid transfer at MCS depends on specific lipid transfer proteins and calcium signaling, operating in concert with lipid metabolic pathways to preserve the unique lipid composition of each organelle membrane [6, 7]. Among these, contacts between the cortical ER (cER) and the PM are particularly prominent, serving as key hubs for the transfer and distribution of phospholipids, sterols, and other lipid species to the PM [8–10]. Proper lipid partitioning in both the PM and endomembranes is essential for the function, trafficking, and turnover of various membrane proteins, including enzymes, transporters, and receptors [11–13]. Notably, ER-PM contact site proteins also influence ER-selective autophagy (ER-phagy) under conditions such as nutrient starvation or ER stress caused by misfolded protein accumulation, highlighting the broader significance of MCS in maintaining ER and cellular homeostasis [14].

Proteins belonging to three distinct families have been shown to be essential elements of ER-PM contact formation and functioning in yeasts, plants and mammals. These proteins, schematically depicted in **Figure 1A**, are: the Ist2/TMEM16/anoctamin/protein family, which includes ion channels and phospholipid scramblases[8, 15], the tricalbin/E-Syts protein family [2, 8, 16, 17], and the VAMP (Vesicle Associated Membrane Protein)-associated protein family, known as VAPs [8, 18, 19]. Here, for simplification, these three types of proteins are referred as Ist2, tricalbin and VAP.

**Figure 1.**
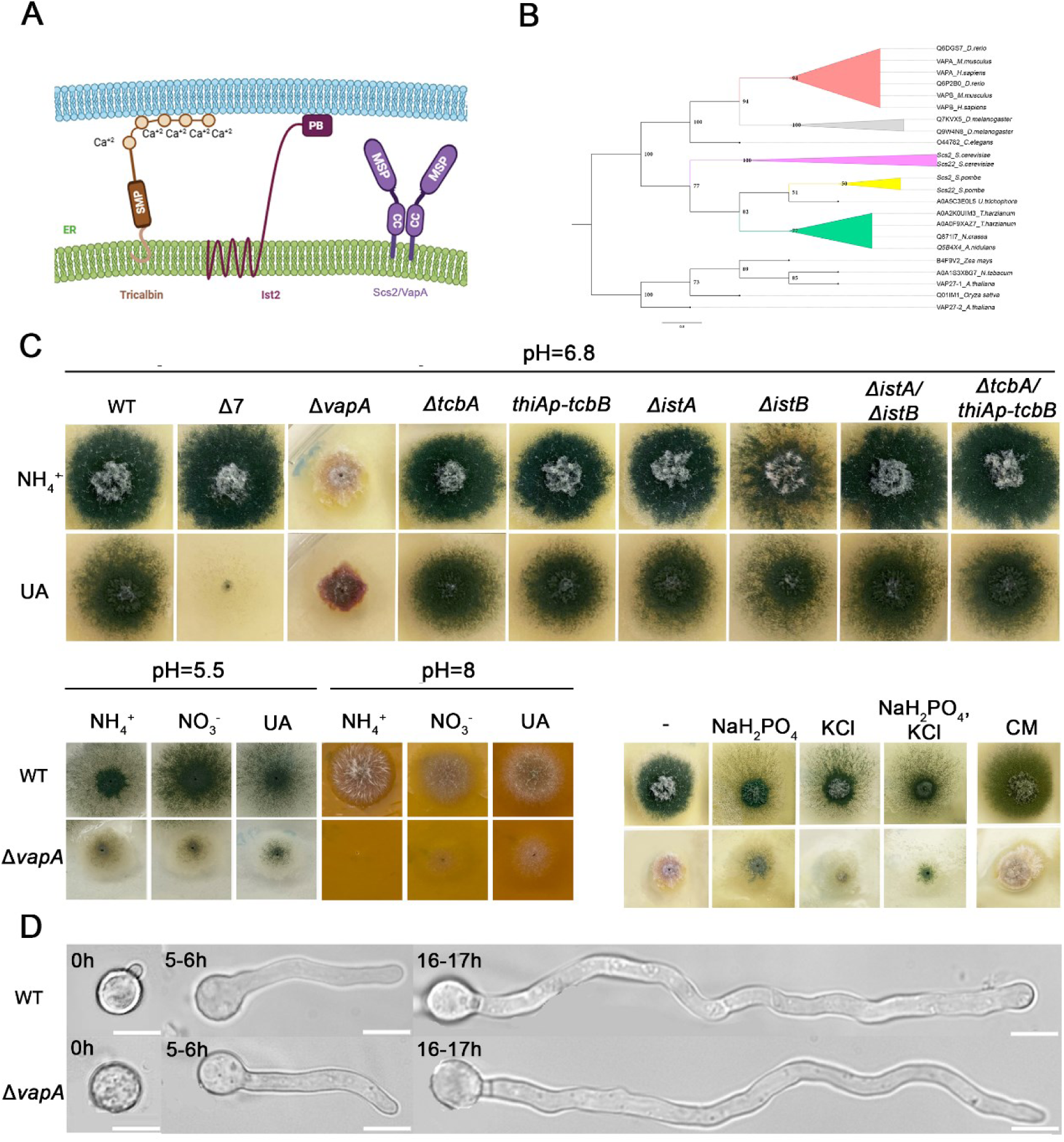
Phenotypic analysis of proteins tentatively associated to ER-PM contacts in *A. nidulans*. **A.** Schema illustrating three families of integral ER proteins that tether the cortical ER to the PM, as characterized in yeasts (*S. cerevisiae* and *S. pombe*) and other eukaryotes: tricalbins, Ist2 and Scs2/VAP proteins. **B.** Phylogenetic relationships of VapA/VAP homologues across model organisms. The tree was reconstructed using the maximum-likelihood method in IQ-TREE, with bootstrap support values shown in black. Organisms are grouped into three major phylogenetic clades: plants (root), fungi and metazoa. The fungal clade diverges from the metazoan lineage, with *S. cerevisiae* forming a distinct branch from the other fungal species. The dataset includes representative model organisms such as *Homo sapiens, Mus musculus, Arabidopsis thaliana, S. cerevisiae, S. pombe, Drosophila melanogaster* and *Caenorhabditis elegans.* The branch scale represents 0.8 substitutions per site, and branch lengths indicate the expected number of substitutions per site. Bootstrap support values were calculated using Ultrafast Bootstrap Approximation (1000 replicates) and SH-aLRT branch test (1000 replicates). **C.** Growth phenotypes of strains carrying single total deletions of genes encoding VapA/VAP, tricalbins or Ist2 proteins (Δ*vapA*, Δ*tcbA*, Δ*istA* and Δ*istB*) or a strain where *tcbB* can be tightly repressed via the *thiA* promoter (*thiA_p_-tcbB*) (**Supplementary Figure S1A**). Growth of doubly knockout/knockdown mutants *ΔistA/ΔistB* and *ΔtcbA/thiAp-tcbB* is also shown. A standard isogenic wild-type strain (wt) and a strain carrying multiple deletions in genes encoding purine/pyrimidine-related transporters are used as controls (Δ*furD* Δ*furA* Δ*fcyB* Δ*uapA* Δ*uapC* Δ*azgA* Δ*cntA*, named Δ7). In the upper panel, strains are grown on minimal media with selected nitrogen sources (ammonium/NH_4_^+^ or uric acid/UA), at pH=6.8, 37°C. Notice that solely Δ*vapA* exhibits dramatically reduced growth rate, conidiospore production (i.e., absence of green color associated with conidiospores) and altered colony morphology. The lower panels depict growth phenotypes at pH=5.5 or pH=8, and on various hyperosmotic media or standard Complete Medium (CM). Notice that the sporulation of Δ*vapA* is somehow enhanced at pH=5.5, and with the addition of NaH_2_PO_4_ and/or KCL in the minimal media. **D**. Microscopy imaging of Δ*vapA,* compared to its isogenic wt strain, at different development stages after germination (0-17 h or germination at 25 ° C). Notice that Δ*vapA* exhibits apparently normal growth rate and germling formation in the time course of microscopy.

In *Saccharomyces cerevisiae*, nearly half of the plasma membrane (PM) lies in close proximity (15–30 nm) to the cortical endoplasmic reticulum (cER), providing a valuable model for studying the formation, function, and physiological roles of ER–PM contact sites and their associated proteins [6, 8, 10, 20–22]. Among these proteins, Ist2, embedded in the ER membrane via eight transmembrane segments, has a long cytoplasmic C-terminal tail that binds PM lipids, thereby tethering the ER to the PM [8, 15]. Loss of Ist2 reduces cER levels, whereas its overexpression enhances ER–PM membrane contact sites (MCS) [8, 23]. The tricalbin proteins (Tcb1/2/3), yeast orthologs of mammalian extended synaptotagmins (E-Syts) and plant synaptotagmins (SYTs) [2, 16, 24], are anchored to the ER via a hairpin loop [16]. These proteins contain a synaptotagmin-like, mitochondrial, and lipid-binding protein (SMP) domain that facilitates lipid binding and transfer [25–27], along with a variable number of C-terminal C2 domains that mediate phospholipid binding and ER–PM tethering [28, 29]. Despite their established role as ER–PM tethers, deletion of tricalbins does not markedly affect cER abundance [8, 30], suggesting their primary function may lie in membrane remodeling or lipid dynamics rather than mechanical tethering. Lastly, Scs2 and Scs22, the yeast homologs of mammalian VAP proteins, are anchored to the ER via a single C-terminal transmembrane segment and contain an N-terminal major sperm protein (MSP) domain. These proteins tether the ER to the PM through interactions between the MSP domain and PM proteins bearing FFAT-like motifs [8, 18]. Deletion of Scs2/22 leads to a significant reduction in cER levels [8]. Each of these tether families contributes uniquely to cER morphology: Scs2/22 are associated with cER sheet formation, while tricalbins promote tubular cER structures at regions of high membrane curvature. While single deletions of Ist2, Scs2/22, or Tcb1/2/3 do not cause major cellular or phenotypic defects, simultaneous deletion of all three tethering systems significantly reduces ER–PM associations and contact site number. This strain, commonly referred to as ‘Δtether,’ exhibits markedly diminished cER and disrupted PM lipid homeostasis [8, 10]. Notably, however, even in the Δtether strain, residual ER–PM contacts persist, supporting that additional factors, as for example the integral ER membrane protein Ice2, contribute to ER–PM association.

ER–PM contact sites, and particularly the role of Scs2/VAP homologues, have also been investigated in the fission yeast *Schizosaccharomyces pombe* [31]. One study demonstrated that ER–PM contacts serve as physical barriers to vesicular secretion, thereby confining exocytosis to the cell tips. This spatial restriction is essential for establishing polarized protein and lipid trafficking required for proper *S. pombe* growth [32]. Another study revealed that eisosome-associated plasma membrane invaginations stabilize local ER–PM contacts through the interaction of Scs2 with Pil1, a core eisosomal component. This interaction influences the remodeling and dynamics of the cortical ER (cER) [33]. Additionally, the same research group showed that Scs2/VAP interactions with phospholipids regulate ER–PM contact formation, and provided evidence that direct binding between Scs2/VAP and Pma1, the major plasma membrane H^+^-ATPase, is crucial for maintaining pH homeostasis [34].

Similar to yeasts, filamentous fungi possess an extensive cER network that forms multiple contacts with the plasma membrane. To date, the only ER–PM tethering protein studied in a filamentous fungus is MoSCS2, a VAP homolog from the rice blast fungus *Magnaporthe oryzae.* Deletion of the MoSCS2 gene results in markedly reduced vegetative growth and conidiospore production, which is associated with decreased virulence [35].

To explore the functional significance of ER–plasma membrane (ER–PM) contact sites in filamentous fungi, we identified and genetically characterized all *Aspergillus nidulans* proteins homologous to Snc2/VAP, Ist2, or tricalbins. We show that the sole Scs2/VAP homolog, VapA, is essential for growth, and provide evidence that this requirement stems from a failure to maintain the polarized localization of specific cargoes at the hyphal apex. This defect correlates with depolarization of the AP-2 adaptor complex, which is required for apical cargo endocytosis [36], and with disruptions in lipid homeostasis. Importantly, VapA deletion does not impair the trafficking, localization, or endocytosis of non-polarized nutrient transporters, suggesting that apical and subapical membrane cargoes depend differentially on partitioning into distinct lipid domains.

## RESULTS

### The VapA/Scs2 homologue is essential for sustaining polarized growth in *A. nidulans*

Most *Aspergillus* species possess homologues of ER–plasma membrane (ER–PM) tethering proteins with significant sequence similarity to those found in *S. cerevisiae* and *S. pombe*. These homologues share approximately 35% identity with *S. cerevisiae* and *S. pombe* proteins, 45% with other filamentous ascomycetes, 41% with basidiomycetes, and 29–30% with mammalian, insect, and plant VAPs. To identify *A. nidulans* homologues of Snc2/VAP, tricalbins, and Ist2 (**Figure 1A**), we performed BLASTp and BLASTx searches using the corresponding *S. cerevisiae* and *S. pombe* proteins as queries. This analysis revealed that *A. nidulans* encodes a single Scs2/VAP homologue (AN4406, hereafter VapA), two tricalbin homologues (AN9149, TcbA; and AN5624, TcbB), and two Ist2 homologues (**AN2477, IstA**; and AN7165, IstB). Using standard reverse genetics based on homologous recombination of DNA cassettes, we constructed null mutants for *vapA, tcbA, istA* and *istB (*Δ*vapA*, Δ*tcbA*, Δ*istA*, and Δ*istB*, respectively). As we were unable to generate a full knockout of tcbB, we instead generated a knockdown strain by replacing the native promoter with the tightly repressible thiA promoter (*thiAp-tcbB*) (**Supplementary Figure S1B)**, as described previously [37, 38].

Phenotypic analysis of these mutants, summarized in **Figure 1C**, revealed that only the Δ*vapA* strain displayed a severe growth defect and a drastic reduction in conidiospore production. To assess potential redundancy, we also generated the double mutants Δ*istA*/Δ*istB* and Δ*tcbA*/*thiAp-tcbB.* Neither of these combinations showed any apparent growth or morphological defects. Standard growth assays were performed on minimal medium (MM) at pH 6.8 and 37°C. We further extended the growth analysis across a range of pH values, temperatures (25, 37, and 42°C), and on complete medium (CM). Under all conditions tested, the severe growth defect remained specific to the Δ*vapA* strain (**Figure 1C** and **Supplementary Figure S1A, Supplementary Figure S2**). Interestingly, the Δ*vapA* defect was modestly alleviated, particularly with respect to conidiospore production, at pH 5.5 or in salt-buffered media. Brightfield microscopy of Δ*vapA* and the isogenic wild-type strain during germination at 25°C showed that Δ*vapA* exhibits normal germling formation and apical growth within the first 17 hours. This suggests that the severe growth impairment observed after 2–4 days of colony development arises from a cellular defect that accumulates at later stages of mycelial maturation and the onset of asexual sporulation. Finally, **Figure 1B** presents a phylogenetic tree of VapA and its homologues across various model organisms. The topology reflects expected evolutionary relationships, with VapA clustering according to major taxonomic lineages such as fungi, mammals, insects, and plants.

### VapA is a cER protein

Since deletion of **VapA**, unlike any other putative ER–PM tether studied, resulted in a severe growth defect, we focused on characterizing its subcellular localization and functional role in more detail. To determine VapA localization *in vivo*, we generated GFP-tagged versions of the protein. Given that both N- and C-terminal regions of VAP proteins may influence their localization and function (see **Figure 1A**), we constructed three variants: GFP fused to the N-terminus (GFP-VapA), the C-terminus (VapA-GFP), and an internal insertion of GFP between the transmembrane (TM) domain, required for ER anchoring, and the functionally important coiled-coil domain (VapA-GFP-TM). To assess the role of ER anchoring in VapA function, we also generated a truncated version lacking the TM domain, with GFP fused to the C-terminus (VapAΔTM-GFP). All constructs were integrated into the endogenous *vapA* locus via homologous recombination, and transformants with correct gene replacement were verified by PCR (see *Materials and Methods* for details). In each case, expression was driven by the native *vapA* promoter to preserve physiological regulation.

All GFP-tagged VapA constructs failed to complement the growth defect of the Δ*vapA* null mutant when expressed under the native promoter (**Figure 2A**, middle panels). However, widefield fluorescence imaging revealed differences in localization depending on the site of GFP insertion (**Figure 2A**, lower panels). Notably, the N-terminally tagged GFP-VapA localized to a membranous network consistent with the endoplasmic reticulum (ER) morphology in *A. nidulans*, primarily labeling cortical membrane patches. In contrast, other constructs, such as VapAΔTM-GFP and VapA-GFP-TM, displayed diffuse cytoplasmic fluorescence with punctate foci suggestive of membrane aggregates, while VapA-GFP exhibited only weak fluorescence.

**Figure 2.**
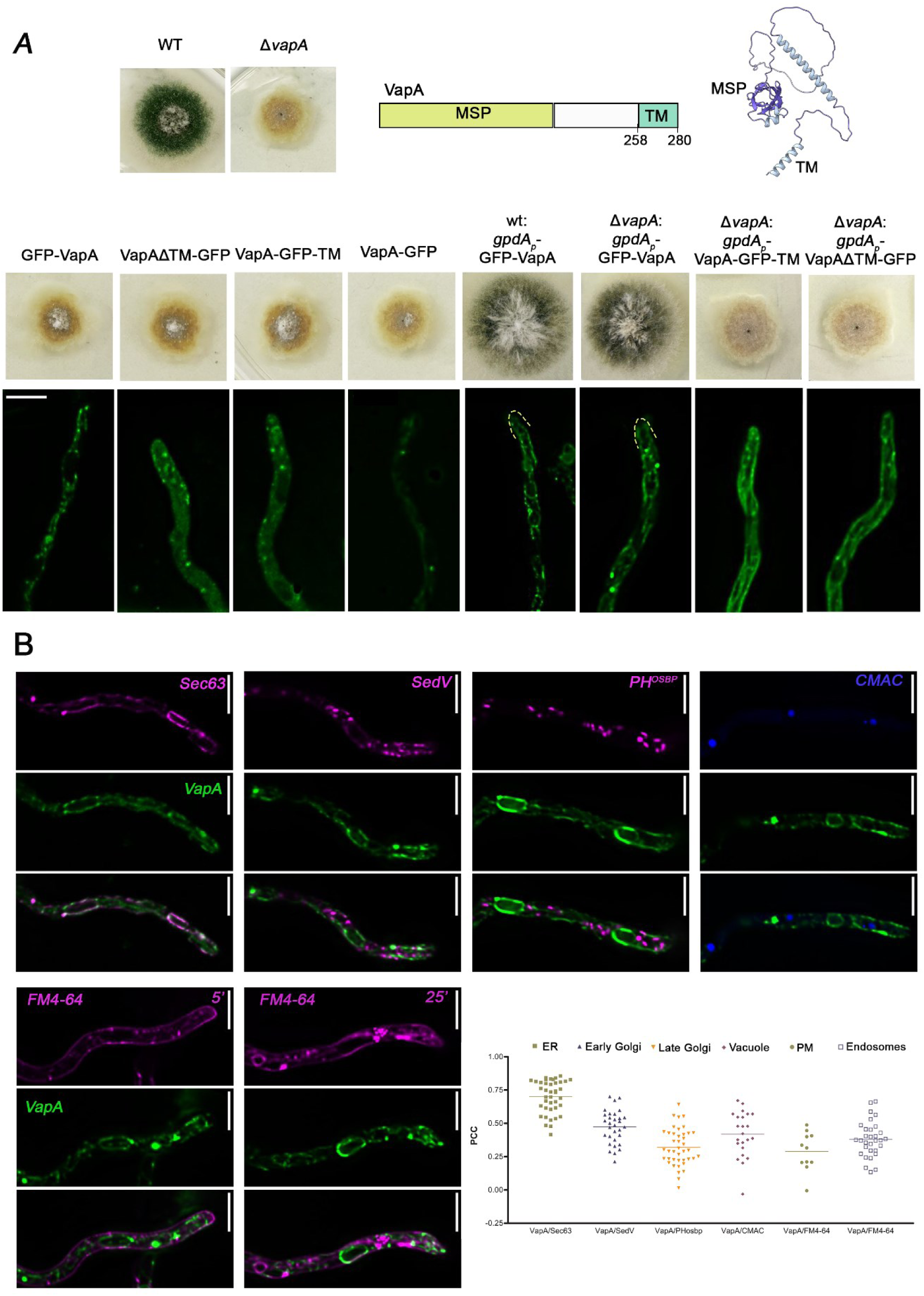
VapA is associated with the cER. **A.** Growth tests of transformants expressing GFP-tagged versions of *vapA* gene. Notice that the *in-locus* substitution of the *vapA* gene with any of the GFP-tagged constructs made (GFP-VapA, VapA-GFP, VapA-ΔΤΜ-GFP, VapA-GFP-TM) did not complement the defective growth phenotype of the original Δ*vapA* recipient strain. Nevertheless, overexpression of GFP-VapA under the *gpdA* strong promoter, unlike that of other constructs, rescued the Δ*vapA* phenotype, indicating that the N-terminal addition of GFP in VapA is functional. On the upper left, there is a cartoon depicting the different domains of VapA, and an alphafold-prediction of the protein (AN4406). Lower panels show maximal intensity projections of deconvolved snapshots indicating the subcellular localization of the corresponding versions of GFP-tagged VapA expressed under the native or the *gpdA* promoter. Notice that uniquely the overexpressed N-terminally tagged GFP-VapA labels a cortical and a perinuclear network resembling ER network. **B**. Colocalization analysis of overexpressed GFP-VapA with standard markers of subcellular compartment (Sec63, SedV, PH^OSBP^, CMAC and FM4-64). Maximal intensity projections of deconvolved z-stacks were used to quantify the co-localization of VapA with these compartment-specific protein markers in live cells. Pearson’s Correlation Coefficient (PCC) values were determined for VapA co-localization with Sec63 (ER; PCC = 0.70 ± 0.12, n = 40), SedV (early Golgi; PCC = 0.47 ± 0.12, n = 32), PH^OSBP^ (late Golgi; PCC = 0.32 ± 0.14, n = 41), FM4-64 for 5 min (endosomes; PCC = 0.38 ± 0.13, n = 31), FM4-64 for 20 min (plasma membrane; PCC = 0.29 ± 0.15, n = 11), and CMAC (vacuole; PCC = 0.42 ± 0.17, n = 23). Each dot represents the PCC of an individual cell. Scale bars: 5 μm

Based on these observations, we hypothesized that the N-terminal GFP-VapA might localize correctly to the ER and retain partial functionality. To test this, and drawing on previous studies in fission yeast indicating that GFP-tagged VapA requires overexpression to restore function in deletion mutants [34], we constructed a version of GFP-VapA driven by the strong constitutive *gpdA* promoter (gpdAp-GFP-VapA). This construct was introduced into both wild-type and Δ*vapA* mutant strains. Expression of *gpdAp*-GFP-VapA successfully complemented the Δ*vapA* growth defect and had no dominant-negative effects in the wild-type background (**Figure 2A**, middle panels). Furthermore, this construct produced strong, ER-compatible fluorescence, consistent with the expected localization of a VAP protein. In contrast, overexpression of other constructs, including VapAΔTM-GFP and VapA-GFP-TM, did not restore function and showed poorly defined, diffuse fluorescence patterns (**Figure 2B**, lower panels). Western blot analysis (**Supplementary Figure S3**) confirmed that only the N-terminally tagged GFP-VapA was stable. Other variants, especially those with internal GFP insertions such as VapA-GFP-TM, were unstable. Based on these results, the gpdAp-GFP-VapA strain was selected for all subsequent analyses.

To precisely identify the compartment labeled by *gpdAp-GFP-VapA* expression, we introduced fluorescent organelle markers into the strain via genetic crossing. These included Sec63 for the endoplasmic reticulum (ER), SedV for the ER-Golgi intermediate compartment (ERGIC) and early-Golgi, and PH^OSBP^ for the late-Golgi and the trans-Golgi network (TGN). Sec63, an essential subunit of the translocase complex, is a well-established ER marker in *A.s nidulans* and other fungi, predominantly labeling the cortical ER (cER) [39]. SedV, a Q/t-SNARE syntaxin required for ER-to-Golgi vesicle transport, localizes to the ERGIC and early Golgi cisternae [38, 40, 41]. PH^OSBP^ is a Pleckstrin Homology (PH) domain that specifically binds phosphatidylinositol-4-phosphate (PI4P), a lipid enriched in the late Golgi and TGN [38, 42, 43]. To further characterize the labeled compartments, we also used the vacuolar lumen marker CMAC [44] and the endosomal/vacuolar membrane dye FM4-64 [45]. As shown in Figure 2B, GFP-VapA strongly colocalizes with the cER marker Sec63 (Pearson’s correlation coefficient, PCC = 0.70), consistent with its role as a VAP protein. Moderate colocalization was observed with the other markers (PCC = 0.29–0.47), reflecting the extensive interactions of the ER with other endomembrane compartments.

### VapA affects the recruitment of enzymes involved in lipid homeostasis at ER–PM contacts

In *S. cerevisiae*, the ER-resident phosphatase Sac1 dephosphorylates PI4P at the plasma membrane (PM) and interacts with the VAP homologues Scs2 and Scs22 to facilitate PI4P turnover specifically at ER–PM contact sites. Loss of Sac1 or its VAP tethers disrupts lipid homeostasis, resulting in PI4P accumulation and altered plasma membrane lipid composition [8, 46]. To determine whether a similar mechanism exists in *A. nidulans*, we constructed a conditional mutant strain in which expression of *sac1* (AN3841) is controlled by the thiamine-repressible *thiA* promoter [47]. As shown in **Figure 3A**, addition of thiamine led to strong repression of *sac1*, severely impairing colony formation and sporulation. These results confirm that *sac1* is essential for viability in *A. nidulans*. To investigate potential functional interactions between Sac1 and VAP proteins, we generated a double mutant strain, Δ*vapA thiAp-sac1*, and assessed its growth phenotype. This strain exhibited a complete absence of colony formation (**Figure 3A**), indicating a strong synthetic negative interaction between VapA and Sac1.

**Figure 3.**
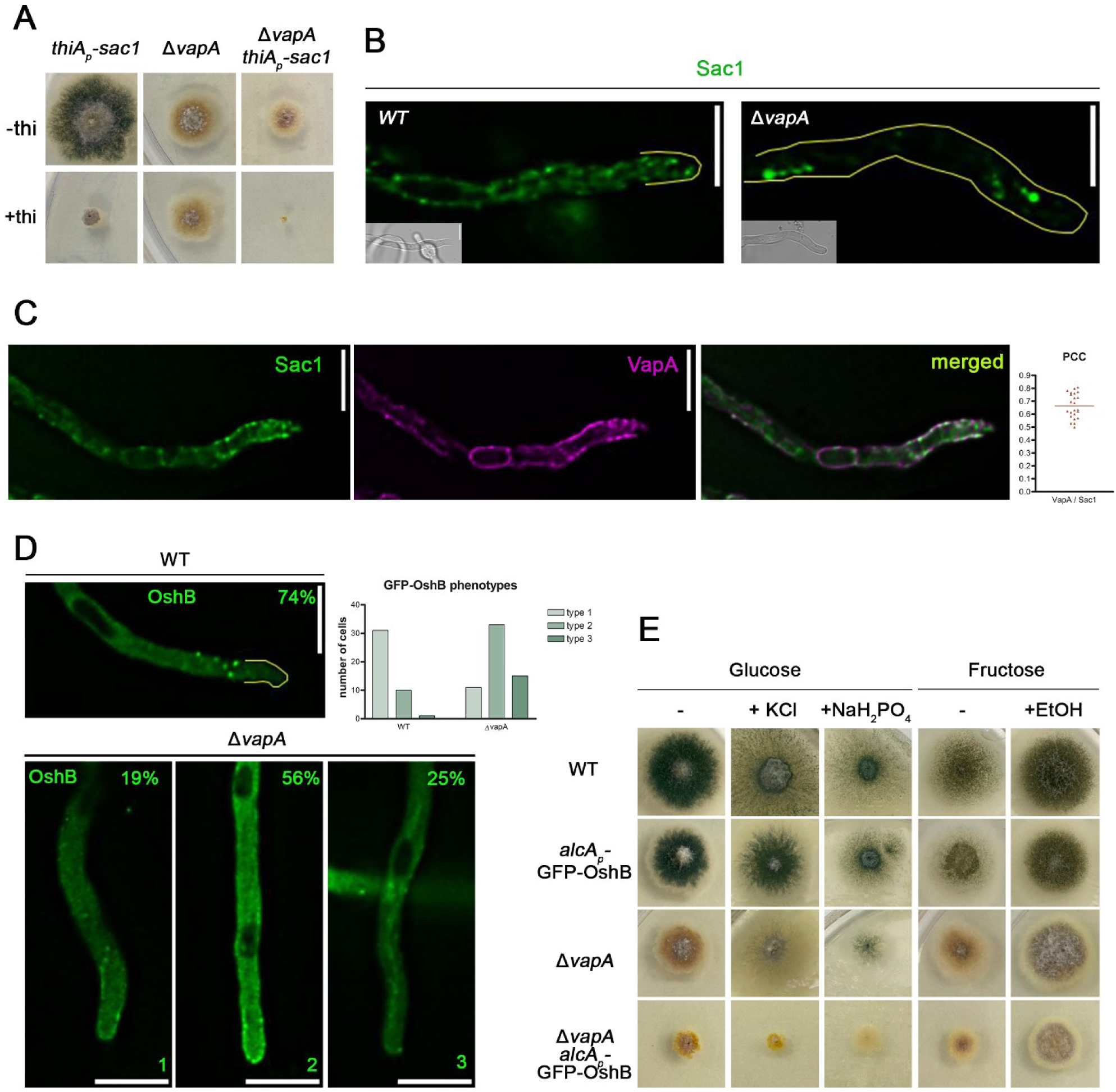
VapA affects the recruitment of Sac1 and OshB at ER–PM Contact Sites. **A.** Growth tests of *thiA_p_-sac1*, Δ*vapA* and Δ*vapA thiA_p_-sac1,* in minimal medium in the presence or absence of thiamine. Notice, *thiA_p_-sac1* has significantly reduced colony size and sporulation, Δ*vapA thiA_p_-sac1* exhibits negative synthetic defect upon thiamine addition. **B.** Maximal intensity projections of deconvolved snap shots showing Sac1-GFP in wt and Δ*vapA* and backgrounds. Notably, the localization of GFP-Sac1 is altered in Δ*vapA.* Sac1 is forming large intracellular aggregates, and displays reduced ER distribution. **C.** Colocalization analysis of mCherry-VapA and Sac1-GFP. Maximal intensity projections of deconvolved z-stacks were used to quantify the co-localization of VapA with Sac1 in live cells. Pearson’s Correlation Coefficient (PCC) values were determined for VapA co-localization with Sac1 (PCC = 0.665 ± 0.096, n = 23). Each dot represents the PCC of an individual cell. **D.** Maximal intensity projections of deconvolved snap shots showing localization of *de novo* expressed GFP-OshB after 4 hours of induction with addition of 0.1% fructose and 0.4% v/v. In Δ*vapA,* three localization patterns were observed: 19% of cells had wt phenotype (type 1), 56% of cells displayed increased cytoplasmic and apical fluorescence (type 2), 25% of cells exhibited diffuse cytosolic haze (type 3). Altogether, GFP-OshB is mislocalized in 81% of Δ*vapA* cells and that percentage is significantly different than the one measured in wt (p<0.05, p = 0.0241, n_wt_=41, n_Δ*vapA*_=59). **E.** Growth tests of strains: wt, *alcA*_p_-GFP-OshB, Δ*vapA and* Δ*vapA alcA*_p_-GFP-OshB, in minimal media containing eighter glucose or fructose as carbon source, and the addition of salts (KCl, NaH_2_PO_4_) or ethanol (EtOH), where indicated. Notice that repression of *oshB* (Glucose containing media) on Δ*vapA* background, results in reduced colony size and complete loss of sporulation. Induction of OshB expression (fructose and ethanol media), partially rescued colony morphology and sporulation of *ΔvapA.* Scale bars: 5 μm.

We next examined the subcellular localization of GFP-tagged Sac1 in both wild-type and Δ*vapA* backgrounds. In wild-type cells, GFP-Sac1 localized to a static, uniformly distributed network adjacent to the plasma membrane, consistent with ER localization. In contrast, in the Δ*vapA* mutant, Sac1 was mislocalized to large intracellular aggregates (**Figure 3B**). These results demonstrate that VapA is required for correct targeting of Sac1 to the ER and ER–PM contact sites.

To assess whether VapA and Sac1 interact *in vivo*, we performed co-localization analysis. GFP-Sac1 exhibited strong co-localization with VapA at cortical ER domains, yielding a Pearson’s correlation coefficient (PCC) of 0.665 (**Figure 3C**). These results support a physical or spatial interaction between VapA and Sac1 at ER–PM contact sites.

Oxysterol-binding homology (Osh) proteins are known regulators of lipid homeostasis and Sac1 activity at ER–PM contacts. In *S. cerevisiae*, Osh3, a PH domain–containing member of this family, functions as a PI4P sensor and Sac1 activator. Importantly, Osh3 recruitment to cortical ER depends on Scs2/Scs22, highlighting a coordinated functional network between VAPs, Osh proteins, and Sac1 [43, 46, 48]. Based on the mislocalization of Sac1 in Δ*vapA* cells, we hypothesized that the localization of OshB, the *A. nidulans* orthologue of Osh3 [49], might also depend on VapA. To test this, we crossed the Δ*vapA* mutant with a strain expressing GFP-OshB under the inducible *alcAp* promoter [49] and analyzed localization in wild-type and Δ*vapA* backgrounds. In wild-type cells, GFP-OshB showed diffuse cytoplasmic fluorescence and distinct cortical foci, particularly at the apical region of hyphae, consistent with its presence at ER–PM contact sites (**Figure 3D** and [49]). In Δ*vapA* cells, OshB localization was disrupted and could be classified into three distinct phenotypes: type 1 (wild-type-like localization, 19% of cells), type 2 (increased cytoplasmic and apical cortical fluorescence, 56%), and type 3 (diffuse cytosolic haze with no clear localization, 25%). Thus, in 81% of Δ*vapA* cells, OshB fails to properly localize to ER–PM contact sites.

To further investigate functional interactions between VapA and OshB, we analyzed the growth phenotype of the Δ*vapA alcAp-GFP-OshB* strain under conditions of *oshB* repression. As shown in **Figure 3E**, repression of *oshB* led to reduced colony size and a complete loss of sporulation. Notably, overexpression of GFP-OshB, achieved by derepression in ethanol/fructose-containing medium, partially restored colony morphology and sporulation in the Δ*vapA* background. These findings suggest that OshB can functionally compensate, at least in part, for the loss of VapA.

Collectively, our results demonstrate that VapA is required for the proper localization and function of both Sac1 and OshB at ER–PM contact sites in *A. nidulans*. These findings support the existence of a conserved tripartite module comprising VAP proteins, ORPs, and PI phosphatases that is essential for plasma membrane lipid homeostasis and fungal growth.

### Deletion of VapA alters PH domain labeling and suggests a role in phospholipid partitioning

We investigated whether deletion of *vapA* affects the subcellular organization and morphology of key organelles, which could underlie the observed growth defects. To this end, we introduced fluorescent molecular markers into the Δ*vapA* mutant background to label the cortical ER (cER; Sec63), ERGIC/early-Golgi (SedV), late-Golgi/trans-Golgi network (PH^OSBP^), nuclei (histone H1), and peroxisomes (AKL-mRFP) [39, 40, 42, 50–52]. In addition to Sec63, we used TcbA-GFP, a tricalbin protein identified in this study, as a marker that more specifically labels cER–plasma membrane contact sites. AKL-mRFP is a chimeric construct that targets mRFP to peroxisomes [51]. As shown in **Figure 4A**, deletion of *vapA* has only minor effects on the morphology of the cER and early Golgi, and no apparent impact on nuclei or peroxisomes. However, we observed a significant 2.5-fold increase in PH^OSBP^ fluorescence at the apical region, suggesting enhanced interaction with phosphatidylinositol 4-phosphate (PI4P). This implies a local accumulation of PI4P at the late-Golgi/TGN in the absence of VapA.

**Figure 4.**
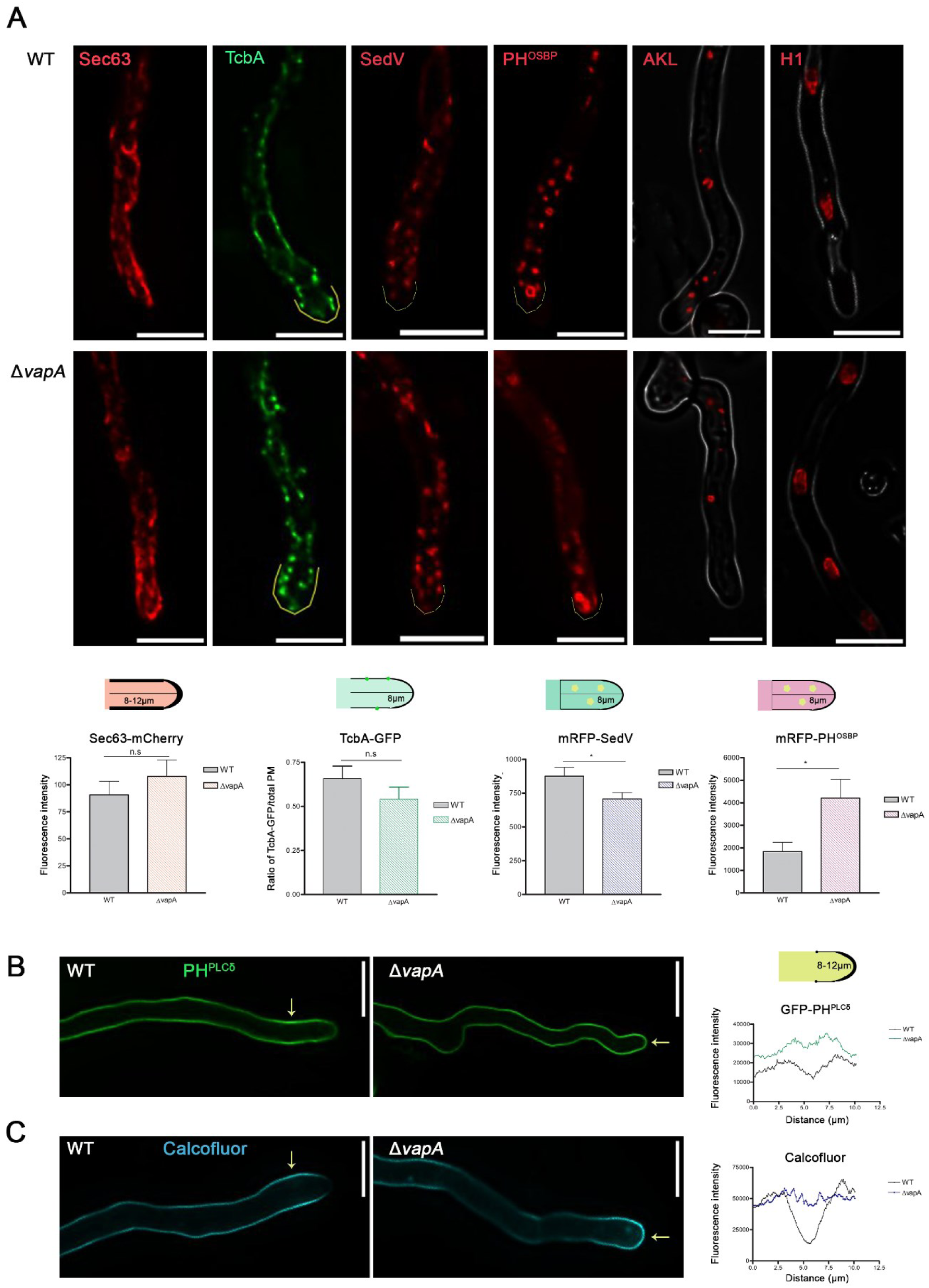
Deletion of vapA modifies labeling by PH and apical chitin deposition. **A.** <u>Upper panel</u>: Maximal intensity projections of deconvolved snap shots showing GFP/mRFP-tagged markers of the ER/cER (Sec63), cER-PM contacts (TcbA), ERGIC/early-Golgi (SedV), late-Golgi/TGN (PH^OSBP^), peroxisomes (AKL-mRFP) and nuclei (H1 histone) in WT and Δ*vapA* backgrounds. <u>Lower panel</u>: Bar plots for the analysis of Sec63, TcbA, SedV and PH^OSBP^ fluorescence respectively. A cartoon on top of each graph illustrates area of the hyphae that has been measured, error bars indicate the mean ± standard deviation for each group. For the statistical analysis of Sec63, TcbA, SedV and PH^OSBP^, an unpaired t test was employed. For Sec63 and TcbA, the test revealed no significant difference in fluorescence intensity between WT and Δ*vapA* groups (WT_Sec63_: Mean ± SEM= 90.63 ± 12.64 N=13, Δ*vapA*_Sec63_: Mean ± SEM= 107.7 ± 15.29 N=17, t=0.8263 df=28, p= 0.4156), (WT_TcbA_: Mean ± SEM= 0.6575 ± 0.07126 N=17, Δ*vapA*_TcbA_: Mean ± SEM= 0.5415 ± 0.06840 N=20, t=1.171 df=35, p= 0.2495). For SedV the test indicates a small but statistically noticeable difference, with Δ*vapA* cells exhibiting relatively lower fluorescence intensity compared to WT (WT_SedV_: Mean ± SEM=875.4 ± 65.84 N=24, Δ*vapA*_SedV_: Mean ± SEM= 706.7 ± 47.21 N=28, t=2.123 df=50, p=0.0387). In PH^OSBP^ there is a significant increase in fluorescence in Δ*vapA* cells compared to WT (Unpaired t test with Welch’s correction: WT_PH_^OSBP^: Mean ± SEM= 1835 ± 409.7 N=20, Δ*vapA*_PH_^OSBP^: Mean ± SEM= 4207 ± 840.0 N=20, t=2.538 df=27, p= 0.0172). **B**.Epifluorescence microscopy showing the PM localization of the PI_4,5_P-lipid marker PH^PLCδ^ in WT and Δ*vapA* background. Notice that PH^PLCδ^ distribution is altered in Δ*vapA*, specifically in the apical PM of hyphae (arrows indicate the difference between the WT and Δ*vapA*). The line plot of GFP-PH^PLCδ^ fluorescence intensity along the PM of the hyphae tip (μm) displays the difference in intensity and distribution of the fluorescent signal (unpaired t test with Welch’s correction: P<0.0001, hyphae measured N_WT_= 27, N_ΔvapA_=30). **C**. Epifluorescence microscopy showing Calcofluor staining in WT and Δ*vapA* background. Calcofluor is a dye staining chitin, an essential component of the fungal cell of wall. Notice that, in Δ*vapA* there is increased chitin deposition in the extreme apex, rather than in the sub-apical collar. Arrows indicate the aforementioned regions in the mutant and WT strain respectively. The line plot of fluorescence intensity along the hyphae tip (μm) highlights the difference in calcofluor staining (unpaired t test with Welch’s correction: P<0.0001, hyphae measured N_WT_= 19, N_ΔvapA_=19). Scale bars: 5 μm. (More details on Materials and Methods section)

To further explore phospholipid distribution at the plasma membrane, we examined a distinct PH domain protein, PH^PLCδ^, which binds specifically to phosphatidylinositol 4,5-bisphosphate (PIP2 or PI(4,5)P_2_) [42, 53, 54]. As shown in **Figure 4B**, loss of VapA resulted in a marked increase in PH^PLCδ^ labeling at the apex of growing hyphae, indicating elevated levels of PI(4,5)P_2_ in this region compared to the wild-type strain. Additionally, calcofluor white staining (Figure 4C) revealed increased fluorescence at the hyphal apex in the Δ*vapA* mutant, suggesting an increase in chitin deposition and altered cell wall composition.

Overall, these results indicate that VapA is required for proper phospholipid partitioning in both the late-Golgi and the apical plasma membrane. Its absence leads to the local accumulation of PI4P and PI(4,5)P_2_, as evidenced by enhanced binding of PH domain probes. These phospholipid changes are also associated with increased chitin deposition at the apex, further underscoring the role of VapA in maintaining phospholipid homeostasis, likely through its functional interactions with Sac1 and OshB.

### VapA is essential for polarized maintenance of growth-related cargoes via its essentiality for apical localization of the AP-2 adaptor complex

The absence of VapA has a dramatic negative effect on hyphal growth, despite having no discernible impact on early developmental stages such as conidiospore germination or polarity establishment. This suggests that VapA may play a critical role in sustaining the maintainance of polarized localization of proteins required for the synthesis and/or deposition of new plasma membrane (PM) and cell wall materials at expanding hyphal tips. To explore this possibility, we examined the steady-state subcellular localization of several key apical proteins: the lipid flippases DnfA and DnfB [36, 55], chitin synthase ChsB [56], and the Q/v-SNARE SynA [69]. Deletion of *dnfA*, *dnfB,* or *chsB* causes severe growth defects and a significant reduction in conidiospore production, phenotypes that closely resemble those observed in the Δ*vapA* mutant, particularly at 25 °C (see **Figure 5A**). Deletion of *synA* results in known to result in only mild growth impairment [69]. In wild-type (wt) cells, these proteins are trafficked to the apical PM via the conventional Golgi- and exocyst-dependent pathway and remain restricted to the apical 2-3 μm region through continuous AP-2-dependent endocytosis and recycling (see schematic in Figure 5A). Although AP-2 is typically considered a clathrin adaptor, we previously showed that in *A. nidulans*, it mediates apical cargo endocytosis independently of clathrin [36]. Similar clathrin-independent roles for AP-2 have also been reported in mammalian central synapses [57]. Notably, in *A. nidulans* deletion of any essential AP-2 subunit causes a severe growth defect, comparable to that seen in mutants lacking DnfA, DnfB, or ChsB [37] (see Figure 5A). Based on these findings, we hypothesized that VapA may be required to maintain the polarized apical localization of these essential cargoes. Because the localization of these proteins shows varying dependence on AP-2, we compared their distribution in Δ*vapA* and Δ*ap2^σ^* mutants and in an isogenic wt control. Strains expressing DnfA, DnfB, ChsB, or SynA in wt and Δ*ap2^σ^*backgrounds were previously described [37], while the corresponding strains in the Δ*vapA* background were generated here through standard genetic crossing (see Materials and Methods).

**Figure 5.**
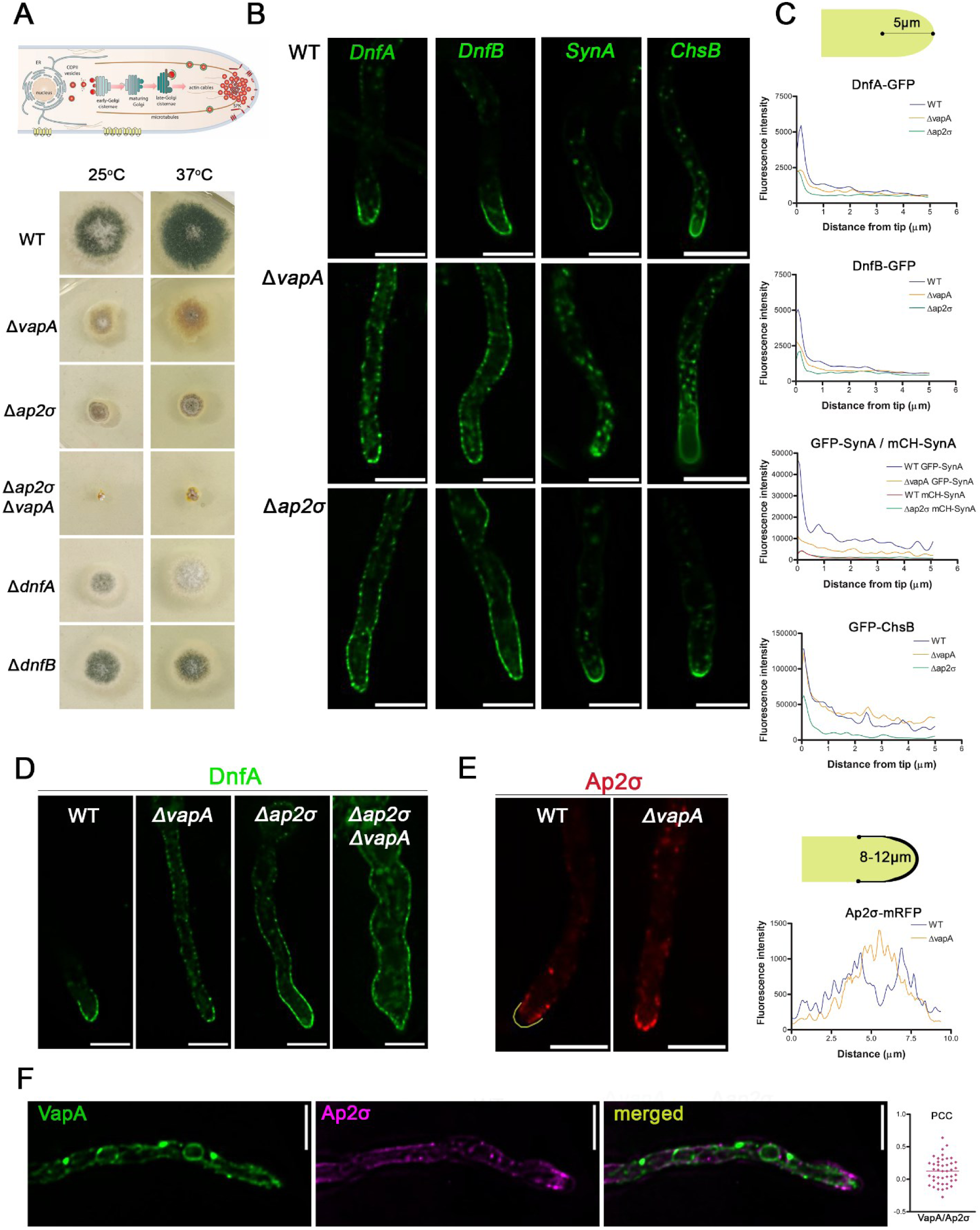
VapA is essential for maintaining the polarized localization of apical membrane proteins and sterol rich domains. **A.** Cartoon depicting the polarized localization of apical markers and growth tests of Δ*vapA*, Δ*ap2^σ^*, Δ*vapA*Δ*ap2^σ^*, Δ*dnfA* and Δ*dnfB* null mutants and an isogenic control strain on standard synthetic minimal medium at 25° and 37° degrees. **B.** Maximal intensity projections of deconvolved snap shots showing the localization of polarized membrane cargoes, DnfA, DnfB, SynA and ChsB in wt, Δ*vapA* and Δ*ap2^σ^* backgrounds. Notice that the apical markers DnfA and DnfB diffused in sub-apical segments of the PM in the Δ*vapA* mutant, rather similarly to Δ*ap2^σ^,* indicating a defect in polarized marker apical maintenance. SynA in Δ*vapA* was largely redistributed to intracellular aggregates, while it remained apically localized in Δ*ap2σ*. ChsB localization appeared little affected in both mutants, yet Δ*ap2σ* exhibited moderately lower fluorescence at the hyphae tip. For better comparison mCherry-SynA Δ*ap2σ* is presented with green in this figure, but the original image and line plot are presented also in **Supplementary figure S4**. Scale bars: 5 μm. **C.** Statistical analysis of cargo localization the three strains. Line plots showing DnfA, DnfB, SynA, ChsB fluorescence intensity along the hyphal tip in wild-type (WT, blue), Δ*vapA* (orange) and Δ*ap2^σ^* (green) strains. The x-axis represents the distance from the hyphal tip (μm), and the y-axis represents fluorescence intensity. WT, Δ*vapA and* Δ*ap2^σ^* exhibit distinct fluorescence distributions along the hyphal axis. To assess spatial distribution differences, the fluorescence profiles were analyzed in 1 µm distance bins from the hyphal tip (0–5 µm). For each bin, pairwise comparisons were conducted and displayed in **Data_Fig5.** Cells measured: DnfA: N_WT_= 18, N_ΔvapA_=22, N_Δap2σ_= 17, DnfB: N_WT_= 26, N_ΔvapA_=28, N_Δap2σ_= 11, SynA: N_WT_=16, N_ΔvapA_=18, N_Δap2σ_= 19, ChsB: N_WT_= 23, N_ΔvapA_=17, N_Δap2σ_= 14. Notice the clear depolarization of DnfA, DnfB and SynA fluorescence from the hyphae apex (0-1 μΜ) in *vapA* null mutant, while in Δ*ap2σ* significant depolarization was solely observed in DnfA and DnfB. **D.** Snap shots showing the subcellular localization of DnfA in WT, Δ*vapA*, Δ*ap2^σ^* and Δ*vapA*Δ*ap2^σ^* backgrounds. In the double mutant Δ*vapA*Δ*ap2^σ^* DnfA displays a rather synthetic phenotype leading to increased mycelium width and increased accumulation of intracellular aggregates. **E.** Epifluorescence microscopy showing the subcellular localization of Ap2^σ^ in wt and Δ*vapA* backgrounds. Notice that Ap2^σ^ seems immobilized on the extreme apex in Δ*vapA*, compared to the WT where Ap2^σ^ mostly marks the sub-apical collar. Line plot shows Ap2^σ^ fluorescence intensity around the hyphal tip, as illustrated in the cartoon, in wild-type (WT, blue) and Δ*vapA* (orange) strains. The x-axis represents the distance from the hyphal tip (μm) and the y-axis represents fluorescence intensity. The fluorescence profiles were analyzed in 1 µm distance bins from 0-10μm. For each bin, pairwise comparisons were conducted and showed significant difference especially between 2-6μm. (P<0.0001, N_WT_= 17, N_ΔvapA_=15). Notice the clear-cut difference at the area of the apex (3-7 μΜ). **F.** Snapshots showing the co-loacalization of Ap2^σ^-mRFP and GFP-VapA, which reveals that Ap2^σ^ and VapA are in close proximity to each other but do not seem to colocalize. Dot plot displays the Pearson’s correlation coefficients (PCC) on y-axis (Mean ± SEM= 0.125± 0.03056, N=43). Scale bars: 5 μm

Our results reveal that deletion of VapA leads to marked depolarization of DnfA, DnfB, and SynA (**Figure 5B and Supplementary figure S4**). In the Δ*vapA* mutant, a significant proportion of these cargoes accumulates in subapical regions of the plasma membrane (PM), often forming a gradient that extends several micrometers from the apical tip into the subapical PM. Noticeably, in Δ*vapA*, SynA also localized to an increased number of cytoplasmic foci. Strikingly, the distribution patterns of DnfA and DnfB in Δ*vapA* closely resemble those observed in the Δ*ap2^σ^* mutant, where the cargoes are similarly mislocalized away from the apical region. Quantitative analysis of fluorescence intensity within 5 μm of the apical tip supported these observations and confirmed a significant reduction in polarized cargo accumulation in the Δ*vapA* mutant (**Figure 5C**). These findings suggest that, as in Δ*ap2^σ^*, cargoes involved in lipid homeostasis in Δ*vapA* fail to undergo efficient apical endocytosis and recycling, resulting in their redistribution to subapical membrane domains. A similar phenotype is observed upon repression of key endocytic genes such as *slaB* and *myoV* (**Supplementary figure S5**). SlaB is an essential early endocytic adaptor that interacts with the AP-2/cargo complex at the PM, while MyoV is a type I myosin motor that functions during the later stages of endocytosis.

To investigate the functional relationship between VapA and the AP-2 complex, we also generated a double null mutant (Δ*ap2^σ^* Δ*vapA*) via genetic crossing. We introduced a DnfA-GFP allele into this strain and compared its localization to that in each single mutant. As shown in **Figure 5D**, DnfA-GFP in the double mutant exhibited a non-polarized PM distribution similar to that in the single parental mutants. Interestingly, the Δ*vapA* Δ*ap2^σ^* strain also showed a heightened cytoplasmic signal associated with a membranous network, indicating a partially additive defect in DnfA localization when both VapA and AP-2 are absent. This cumulative disruption in DnfA trafficking correlated with reduced colony growth and pronounced morphological alterations in hyphae (**Figures 5A and 5D**).

To further investigate the functional relationship between VapA and AP-2, we expressed a functional GFP-tagged version of AP-2 in the Δ*vapA* mutant. As shown in **Figure 5E**, the absence of VapA disrupts the normal localization of AP-2-GFP, leading to increased accumulation at the apex and a corresponding reduction in the subapical collar region (2–3 μm from the tip), where active endocytosis and recycling of growth-related cargoes typically occur. Since AP-2 is recruited to the plasma membrane (PM) through a tripartite interaction with cargo proteins and specific phospholipids, particularly phosphatidylinositol 4,5-bisphosphate (PI(4,5)P_2_), these findings suggest that VapA loss alters the organization of PM lipid domains, impairing AP-2 retention in the subapical collar and thus its ability to interact with apical cargoes. The additive effects observed in AP-2 and VapA double mutants further imply that VapA may also influence earlier trafficking events, likely involving anterograde cargo transport. Lastly, to examine whether VapA and AP-2 physically interact, we assessed their spatial distribution. As shown in **Figure 5F**, VapA and AP-2 do not colocalize, indicating that VapA likely affects AP-2 localization indirectly, most plausibly by modulating PM lipid composition.

### Further evidence supporting that VapA modifie*s* membrane lipid partitioning in the PM

As several apical markers are thought to partition within sterol-rich membrane domains in fungi [58], we investigated whether ergosterol distribution is altered in the Δ*vapA* and Δ*ap2^σ^* mutants compared to the wt strain, which typically exhibits enriched filipin staining at hyphal tips [59]. Filipin dye staining revealed that deletion of either VapA or AP-2 resulted in a pronounced depolarization of filipin accumulation, with staining appearing uniformly distributed along the plasma membrane (PM) rather than concentrated at the tips (**Figure 6A**). This suggests that ergosterol partitioning is disrupted in both mutants, leading to a loss of polarized membrane organization. Given this apparent alteration in ergosterol localization, we next assessed whether the Δ*vapA* mutant displayed altered sensitivity to ergosterol-targeting antifungal agents relative to the wt and Δ*ap2^σ^* strains. As shown in **Figure 6B**, the Δ*vapA* mutant was significantly more sensitive to subtoxic concentrations of itraconazole than either the wt or Δ*ap2^σ^* strain. This finding supports the idea that VapA contributes to processes upstream of endocytic cargo trafficking, likely including the regulation of membrane sterol composition.

**Figure 6.**
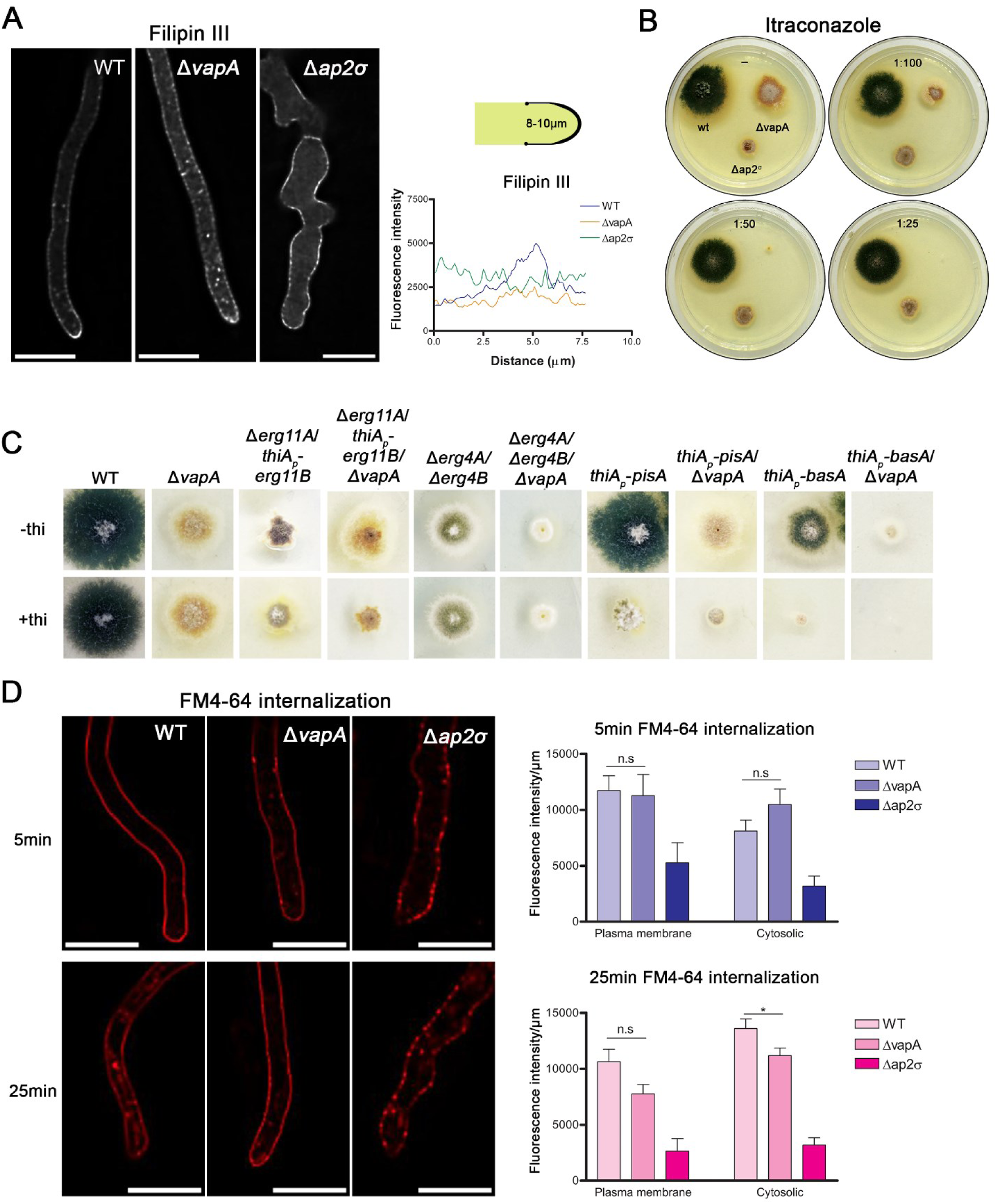
Evidence that VapA is crucial for proper membrane lipid homeostasis and partitioning to the PM. **A.** Snap shots showing filipin III staining of ergosterol in WT, Δ*vapA*, Δ*ap2^σ^*. Notably, the null mutants exhibit distinct staining patterns, indicating altered ergosterol distribution at the plasma membrane and in particular reduced apical localization. This is statistically confirmed in a line plot, which represents filipin III fluorescence intensity along the hyphal tip (μm), as illustrated in the carton on the right. Specifically, to investigate spatial distribution differences, a Kruskal-Wallis test was first applied across the three strains, revealing significant global differences (P<0.0001; H = 152.3, cells measured: N_WT_= 20, N_ΔvapA_=32, N_Δap2σ_= 18). The fluorescence profiles were analyzed in 1 µm distance bins from 0-8μm. For each bin, pairwise comparisons were conducted and showed significant difference. (P<0.0001). **B.** Growth phenotypes of WT, Δ*vapA* and Δ*ap2^σ^* on minimal medium supplemented with subtoxic concentrations of itraconazole. Δ*vapA* displays heightened sensitivity to itraconazole even at low concentrations, in contrast to WT and Δ*ap2σ*, which exhibit normal or near-normal growth. **C.** Growth test of various strains repressed for genes involved in lipid biosynthetic pathways, on minimal medium in the presence or absence of thiamine. Strains include WT, Δ*vapA,* Δ*erg11A/thiAp-erg11B,* Δ*erg11A/thiAp-erg11B/*Δ*vapA,* Δ*erg4A/*Δ*erg4B,* Δ*erg4A/*Δ*erg4B/*Δ*vapA, thiAp-pisA, thiAp-pisA/*Δ*vapA, thiAp-basA, thiAp-basA/*Δ*vapA.* Notably, the combination of Δ*vapA* with repression or deletion of genes involved in lipid biosynthesis - *erg11A/B, erg4A/B,* and *pisA* - results in a synthetic growth defect, indicating potential genetic interactions between VapA and key lipid metabolic pathways. **D.** Maximal intensity projections of deconvolved snap shots showing FM4-64 distribution in WT, Δ*vapA* and Δ*ap2^σ^* at 5min and 25min post-staining. Unpaired t-test between WT and Δ*vapA*, revealed a statistically significant decrease in cytosolic fluorescence intensity in Δ*vapA* at 25 minutes (p= 0.0308, t=2.201 df=75, N_WT_= 38, N_ΔvapA_=39), indicating delayed or altered FM4-64 internalization. Scale bars: 5 μm. For additional interpretation see main relevant text.

To further probe the role of VapA in membrane lipid homeostasis, we tested for synthetic growth defects in double mutants combining Δ*vapA* with alleles affecting major membrane lipid biosynthetic pathways. These included: (a) double knockout or knockdown mutants in ergosterol biosynthesis (Δ*erg11*A/*thiAp-erg11B* and Δ*erg4A*/Δ*erg4B*), (b) a knockdown mutant of phosphatidylinositol synthase (*thiAp-pisA*), and (c) a knockdown mutant of sphingolipid C4 hydroxylase (*thiAp-basA*), as previously described [11]. In all cases, the resulting double mutants exhibited exacerbated growth defects compared to their respective single mutants, indicating that loss of VapA imposes a synthetic burden on strains with impaired lipid biosynthesis (**Figure 6C**).

We also examined the impact of VapA on endocytosis using FM4-64, a lipophilic fluorescent dye that initially incorporates into the PM and is subsequently internalized via bulk endocytosis. FM4-64 internalization is sensitive to PM fluidity and lipid composition. In the Δ*vapA* mutant, initial PM labeling was comparable to the wt after 5 minutes; however, the dye failed to label internal endosomal membranes even after 25 minutes, a time point at which robust internalization is normally observed in the wt (**Figure 6D**). This indicates that PM lipid composition and fluidity are significantly altered in the absence of VapA. Interestingly, FM4-64 uptake was also delayed in the Δ*ap2^σ^* mutant. Moreover, at the 5-minute time point, FM4-64 labeling in Δ*ap2^σ^* was already distinct from both wt and ΔvapA, displaying punctate cortical staining rather than a uniform PM signal, further supporting a functional link between VapA and the AP-2 complex

### VapA is crucial for polarized recruiting of the endocytic machinery at the apical region

Since VapA was shown to influence apical cargo endocytosis by modulating AP-2, we investigated whether it also affects additional key components of the endocytic machinery that function downstream of AP-2, such as SlaB^Sla2/End4^, SagA^End3^, AbpA^Abp1^ or MyoA^Myo5^ [36, 60]. As previously mentioned, SlaB is an adaptor protein that, together with Ent1 and Ent2, forms a midcoat complex following AP-2’s interaction with cargo. SagA is an EH domain-containing protein that forms a complex with Sla1 and Pan1, contributing to the next layer of the endocytic coat and playing a central role in actin organization. AbpA is an essential actin-binding protein, while MyoA represents the major class I myosin motor involved in the late stages of actin-driven endocytosis, functioning independently of earlier endocytic steps [61].

These endocytic proteins typically accumulate in a polarized manner at the subapical collar of *A. nidulans* and other filamentous fungi, a region characterized by high levels of constitutive endocytosis [36, 37, 62–67]. To assess whether VapA affects their localization, we generated functional fluorescent chimeras of SlaB-GFP, SagA, AbpA-dsRED, and MyoA-GFP (Materials and Methods), and introduced them into the Δ*vapA* strain. We then compared their steady-state subcellular localization to that observed in an isogenic wild-type strain and in the Δ*ap2^σ^* mutant background (**Figure 7**). Although some minor differences were observed in the localization of individual proteins across the three strains, both imaging and quantitative analysis revealed a consistent trend: deletion of *vapA* or *ap2^σ^* reduced the polarization of these proteins. Specifically, their accumulation in the subapical collar region (1–3 μm from the apex) decreased, while their presence at the apex (0–1 μm) increased. These findings support the idea that the absence of VapA depolarizes AP-2 and this in turn reduces the recruitment of early endocytic factors in the subapical collar region.

**Figure 7.**
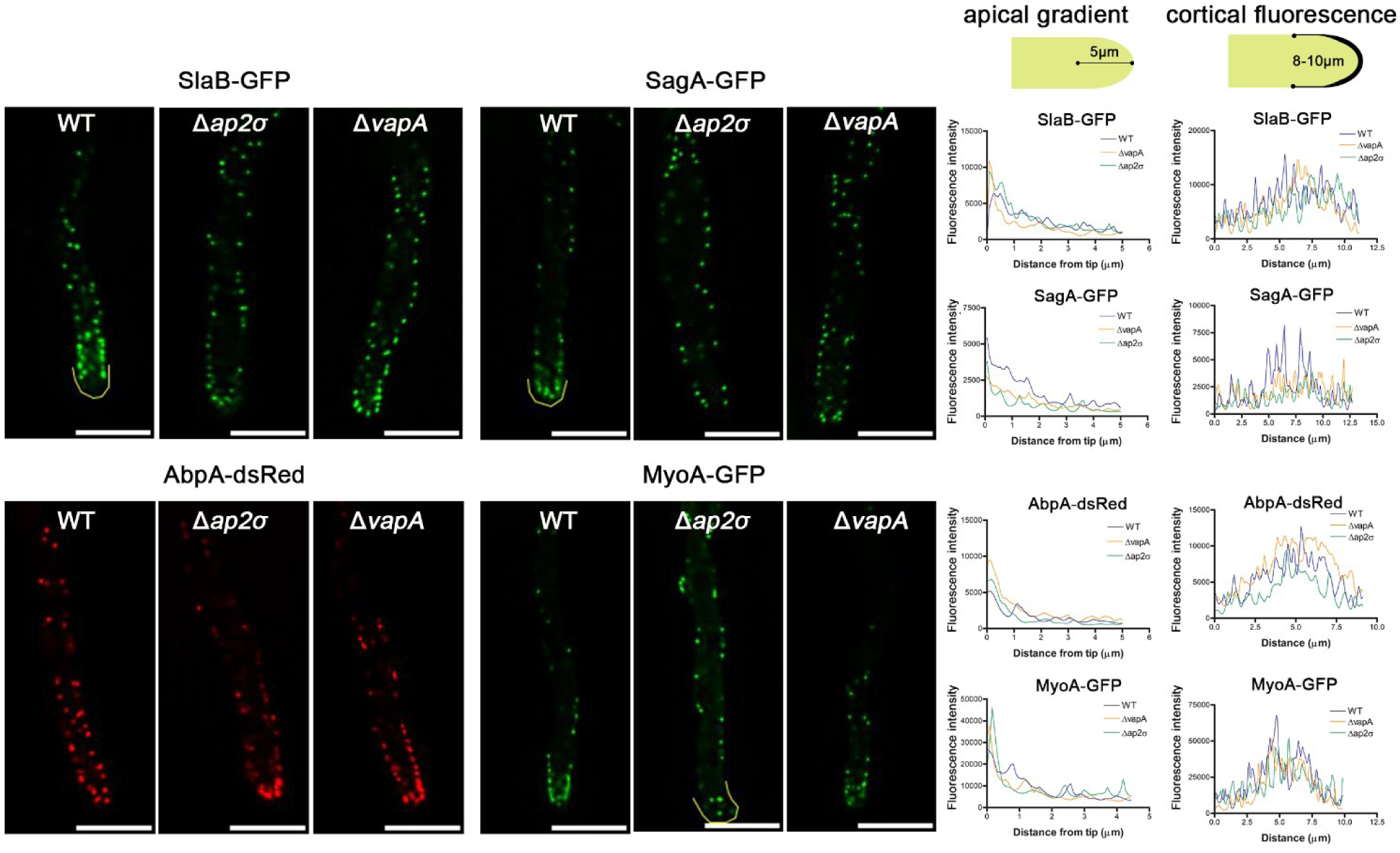
VapA is crucial for the proper functioning of the endocytic machinery at the apical region. **A.** Epifluorescence microscopy showing the subcellular localization of endocytic markers SlaB, SagA, AbpA and MyoA in WT, Δ*ap2^σ^* or Δ*vapA*, backgrounds. SlaB and SagA lost their apical localization in Δ*vapA* and Δ*ap2σ*. AbpA, which marks PM-associated actin patches, showed increased localization at hyphae tips in Δ*vapA* and Δ*ap2^σ^,* compared to WT. MyoA seemed little affected in Δ*vapA,* while it showed increased localization in non-polarized regions of the hyphae in Δ*ap2σ.* Line plots quantifying AbpA, SlaB, SagA, MyoA fluorescence intensity along the hyphal tip (as illustrated in the cartoon) in wild-type (WT, blue) and Δ*vapA* (orange) and Δ*ap2^σ^* (green) strains. The x-axis represents the distance from the hyphal tip (μm), and the y-axis represents fluorescence intensity. To further access different labelling patterns, Curve plots displaying fluorescence intensity around the hyphal tip (as illustrated in the cartoon). The fluorescence profiles were analyzed in 1 µm distance bins, 0–5 µm for line plots, 0-12 µm for curve plots. For each bin, pairwise comparisons were conducted and displayed in **Data_Fig7**. Scale bars: 5 μm

### VapA has a minor role in the secretion of apical cargoes

Apart from its major role in apical cargo endocytosis and recycling, the lack of VapA might also lead to a minor defect in cargo secretion at the apical region. This was suggested by the observation that in several hyphae of the Δ*vapA* or Δ*vapA*Δ*ap2^σ^*mutants we could also detect an increase in labeling of cytoplasmic membrane-like structures, mostly detected by DnfA-GFP. (see **Figures 5B, 5D** and **8**). These cytoplasmic structures might correspond to the presence of cargoes in secretory compartments (e.g., *late*-Golgi/TGN or *post*-Golgi secretory carriers) that fail to reach the apical segment or delayed in their secretion. To investigate this possibility, we recorded the colocalization of DnfA-GFP with fluorescent markers of the *late*-Golgi/TGN (mRFP-PH^OSBP^) and *post*-Golgi carriers (AP-1^σ^-mRFP) [37] in wt and Δ*vapA* backgrounds. **Figure 8** shows that in the absence of *vapA*, DnfA-GFP appears to have increased colocalization with late-Golgi (mRFP-PH^OSBP^) and moderately reduced colocalization with AP-1^σ^-mRFP. These finding reveals that VapA has also a minor role in the dynamics of cargo secretion, which under the light or other results presented here is due its role in phospholipid homeostasis.

**Figure 8.**
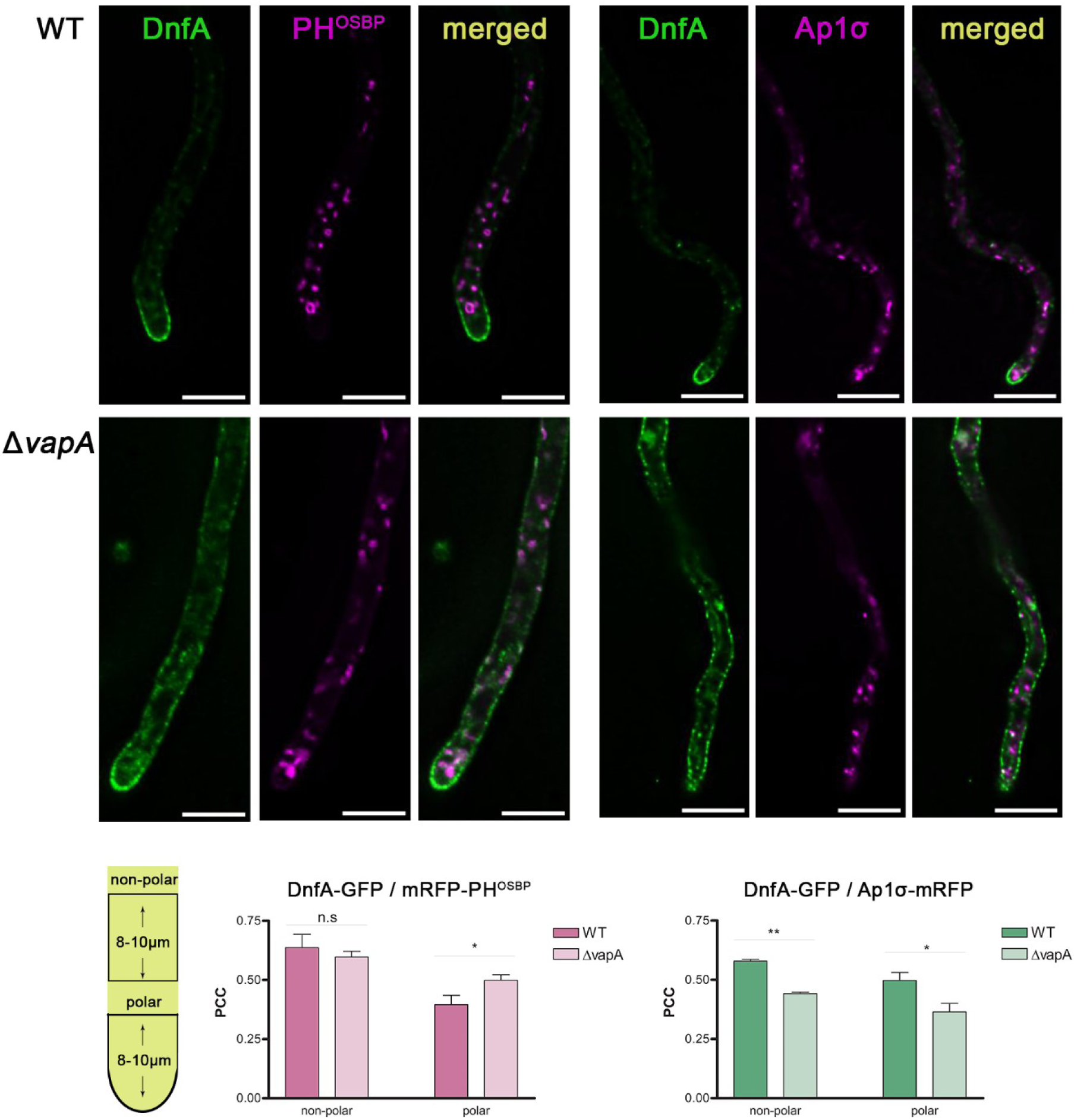
VapA has a minor role in the secretion of apical cargoes. Epifluorescence microscopy showing the co-localization of DnfA with PH^OSBP^, Ap1^σ^ or FM4-64 in wt and Δ*vapA.* Co-localization was assessed in both apical (polarized) and subapical (non-polarized) regions of hyphae, as indicated in the accompanying scheme. There was no significant difference between the wt and *vapA* null mutant in DnfA/FM4-64 co-localization experiments. However, DnfA exhibited significantly reduced overlap with Ap1σ in Δ*vapA* compared to WT in both regions measured (non-polar: PCC_wt_= 0.579, PCC_Δ*vapA*_= 0.442, p_-value_: 0.0055, N_WT_= 20, N_ΔvapA_=32, polar: PCC_wt_= 0.4972, PCC_Δ*vapA*_= 0.36480, p_-value_: 0.0244, N_WT_=16, N_ΔvapA_=32) and increased co-localization with the late Golgi marker PH^OSBP^, specifically at the hypha tip (polar: PCC_wt_= 0.395, PCC_Δ*vapA*_= 0.499, p_-value_: 0.0268). This result might signify a defect or delay in packaging of DfnA in post-Golgi AP-1 secretory vesicles. Accompanying bar plots display Pearson’s correlation coefficients (PCC) on the y-axis, separated by polarized and non-polarized measurements for WT and Δ*vapA.* Scale bars: 5 μm

### VapA is redundant for the localization of transporters and other non-apical cargoes

All evidence obtained strongly suggested that in the Δ*vapA* mutant polarized apical cargoes cannot undergo proper AP-2 dependent endocytosis. In addition, the Δ*vapA* deletion also seems to reduce the steady state accumulation of key endocytic factors at the subapical collar region. In *A. nidulans,* however, there is a distinct mechanism of endocytosis operating at the subapical segments of the PM. This concerns the turnover of several nutrient transporters, which are localized to the PM in a rather antipolar manner (i.e. ‘missing’ from the apical tips; [36, 68]. The endocytosis of these transporters occurs in response to specific physiological or stress signals and is clathrin-dependent, but surprisingly AP-2 independent [36]. Also notably, these transporters translocate to the PM via a Golgi-independent route, distinct from the conventional secretion mechanism of apical cargoes [38, 68–70]. Golgi-independent translocation to the PM is also followed by cargoes other than nutrient transporters, as for example the proton pump ATPase PmaA or the PalI component of the alkaline pH sensing system [68]. Thus, we were interested in addressing whether VapA has also an effect on the localization and/or endocytosis of transporters and other non-polarized cargos.

We first tested whether the genetic deletion of VapA compromises the activity of several transporters by affecting their trafficking to the PM. This can be scored by simple growth tests where we use toxic analogues of substrates transported by well-characterized specific transporters. For example, oxypurinol (OX), 5-fluorouracil (FU), 5-fluorocytosine (FC) or 8-azaguanine (8AZG) are toxic to wt *A. nidulans* due to their specific uptake by the UapA/UapC, FurD, FcyB and AzgA transporters, respectively [71]. A strain carrying multiple deletions of gene encoding these transporter genes (named Δ7) shows increased resistance to these purine analogues [72, 73]. This is demonstrated in **Figure 9A**. In the same growth test, the Δ*vapA* null mutant showed high sensitivity to all toxic analogues tested, suggesting that the respective transporters (i.e., UapA, UapC, FurD, FcyB and AzgA) are active, which in turn means that they translocate normally to the PM.

**Figure 9.**
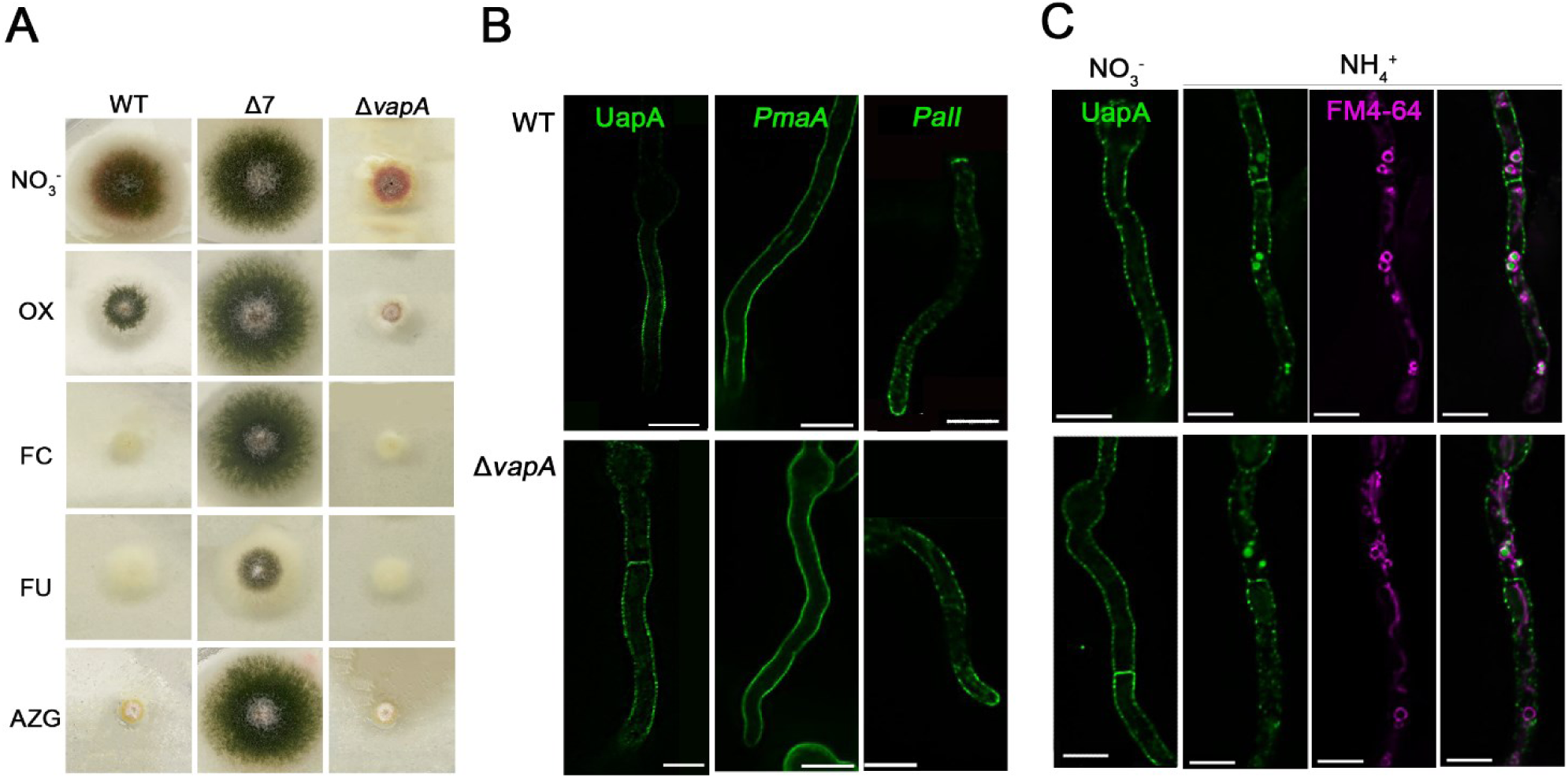
VapA is redundant for the localization and endocytosis of transporters and other non-apical cargoes. **A.** Growth tests of Δ*vapA* in toxic nucleobase analogues. The toxic analogues used, oxypurinol (OX), 5-fluorocytosine (FC), 5-fluorouracil (FU), 8-azaguanine, were added in media containing nitrate as a nitrogen source, at 37°C. A wt strain was used as a negative control and Δ7 as a positive control. These analogues are incorporated in the cells by several different transporters (OX by UapA, FC by FcyB, FU by FurD and AZG by AzgA; The Δ7 strain lacks these transporters, as well as other secondary related transporters, and is thus resistant to the toxic nucleobase analogues; see ref). Δ*vapA* shows the same phenotype as the isogenic wt strain in the presence of toxic nucleobases, strongly suggesting that all major nucleobase transporters are functionally localized to the PM. **B**. Maximal intensity projections of deconvolved snap shots showing the localization of non-polarized membrane cargoes, UapA, PmaA (PM proton pump ATPase) and PalI (pH sensing component of the PM) in wt and Δ*vapA.* The results confirm that all these non-polarized transmembrane cargoes have normal PM distribution in Δ*vapA.* **C**. Epifluorescence microscopy showing the endocytosis of UapA in wt and Δ*vapA.* UapA transporter is derepressed overnight in the presence of nitrate, then incubated with ammonium for 2h, to trigger its internalization and subsequent vacuolar degradation. FM4-64 was used to monitor endosomal and vacuolar compartment. The results showed that UapA is localized in vacuoles in both strains (yellow arrows) after addition of NH4^+^, indicating that transporter endocytosis and turnover is unaffected in Δ*vapA.* Scale bars: 5 μm.

To directly investigate whether the Δ*vapA* deletion has any effect on the trafficking of transporters, we decided to examine the localization of the well-studied UapA purine transporter and two other non-polarized PM cargoes, the essential proton pump ATPase PmaA and the PalI component of an alkaline sensing receptor complex. For achieving this, we constructed by genetic crossing strains expressing functional GFP-tagged versions of these cargoes in the Δ*vapA* background. **Figure 9B** shows that in all cases these transmembrane proteins properly translocate in the PM, similarly to the wt.

As the UapA transporter is a cargo that undergoes regulated endocytosis in response to specific signals, we also tested whether its endocytosis is affected in the genetic absence of VapA. More specifically, we asked whether UapA endocytosis elicited by the addition of ammonium in the growth medium [74] operates properly in the Δ*vapA* mutant. **Figure 9C** confirms that UapA endocytosis takes place properly and is followed by sorting and turnover of UapA in the vacuoles, as in the wt background.

Overall, our results show that unlike the dramatic effect the VapA deletion has on the polarized maintenance of apical cargoes, reflected in growth, defects, the absence of VapA does not affect either translocation to the PM nor the endocytosis of cargoes that tend to be localized to the PM in a non-apical fashion and be endocytosed by an AP-2 independent mechanism. The importance of this finding is discussed further in the next section.

All evidence obtained strongly suggests that in the Δ*vapA* mutant, polarized apical cargoes cannot undergo proper AP-2–dependent endocytosis. Additionally, the deletion of VapA appears to reduce the steady-state accumulation of key endocytic factors at the subapical collar region. However, in *A. nidulans,* a distinct endocytic mechanism operates at the subapical regions of the plasma membrane (PM), specifically involved in the turnover of several nutrient transporters. These transporters are localized in an antipolar manner—i.e., they are absent from the apical tips [36, 68]. Their endocytosis occurs in response to specific physiological or stress signals and, while clathrin-dependent, is surprisingly independent of AP-2 [36]. Notably, these transporters also translocate to the PM via a Golgi-independent route, distinct from the conventional secretion pathway used by apical cargoes [38, 68–70]. This unconventional trafficking route is also utilized by other non-polarized cargoes, such as the proton pump ATPase PmaA and PalI, a component of the alkaline pH-sensing system [68].Given these distinct mechanisms, we sought to determine whether VapA influences the localization and/or endocytosis of nutrient transporters and other non-polarized cargoes.

To assess whether VapA deletion affects transporter activity by impairing trafficking to the PM, we performed growth assays using toxic analogues of transporter substrates. These analogues, oxypurinol (OX), 5-fluorouracil (FU), 5-fluorocytosine (FC), and 8-azaguanine (8AZF), are taken up by the UapA/UapC, FurD, FcyB, and AzgA transporters, respectively [71]. Wild-type *A. nidulans* is sensitive to these compounds due to efficient uptake, whereas a strain carrying deletions of seven relevant transporter genes (Δ7) shows resistance [72, 73], as demonstrated in **Figure 9A**. Under the same conditions, the Δ*vapA* mutant exhibited high sensitivity to all toxic analogues tested, indicating that these transporters are functional and reach the PM normally.

To directly assess PM localization, we examined the distribution of GFP-tagged versions of three representative non-polarized PM cargoes: the purine transporter UapA, the essential ATPase PmaA, and PalI. These constructs were introduced into the Δ*vapA* background via genetic crosses. As shown in **Figure 9B**, all three proteins localized correctly to the PM in the ΔvapA mutant, similarly to wild-type. Because UapA undergoes regulated endocytosis in response to ammonium addition [74], we also investigated whether this process is affected by VapA deletion. **Figure 9C** shows that UapA is internalized and sorted into vacuoles in ΔvapA cells just as in wild-type, indicating that its endocytosis proceeds normally.

In summary, our results demonstrate that while VapA is critical for the polarized, AP-2–dependent maintenance of apical cargoes, it is not required for the Golgi-independent trafficking or AP-2–independent endocytosis of non-polarized PM cargoes such as UapA, PmaA, and PalI. This distinction highlights the compartmentalized nature of endocytic pathways in *A. nidulans* and supports the idea that VapA specifically regulates endocytosis at the apical domain.

## DISCUSSION

The physiological importance of VAPs in eukaryotes is underscored by studies showing that their genetic depletion in mammals is associated with several neurological disorders, including amyotrophic lateral sclerosis (ALS), Alzheimer’s disease (AD), and α-synucleinopathies such as Parkinson’s disease (PD) and multiple system atrophy (MSA) [75–77]. Moreover, mammalian VAPs are hijacked by various viruses and intracellular bacteria to support their replication [13, 78]. In plants, VAPs are also essential, playing key roles in lipid homeostasis, ER function, endocytosis, and interactions with fungi [79, 80].

In fungi, ER–plasma membrane (ER-PM) contact site proteins have been predominantly studied in unicellular yeasts. Here, we genetically characterized all six predicted ER-PM contact site proteins in the model filamentous fungus *A. nidulans*, and found that only the single VAP/Scs2 homologue, VapA, is essential for proper growth and conidiospore formation. A similar growth defect has been observed upon deletion of the VAP/Scs2 orthologue in another filamentous fungus, the rice blast pathogen *M. oryzae* [35], where its loss also reduces virulence toward plant hosts. In contrast, deletion of VAP/Scs2 homologues in the yeasts *S. cerevisiae* and *S.pombe* causes only mild growth defects [8, 31, 46]. This discrepancy likely reflects differences in growth strategies: unlike yeasts, filamentous fungi rely not only on the establishment but also the maintenance of cell polarity, which supports their development of long, asymmetric, coenocytic hyphae rather than single cells [63, 67, 81].

To confirm VapA as an ER-resident protein, we functionally fused GFP to its N-terminus and showed colocalization with the ER marker Sec63, establishing it as an integral ER membrane protein. We further demonstrated that the C-terminal transmembrane (TM) domain is required for both ER anchoring and for rescuing the growth defect of the Δ*vapA* deletion mutant. Notably, as observed in *S. pombe* [34], full complementation of Δ*vapA* by GFP-VapA required high-level expression driven by a strong promoter, suggesting that VAP function is dosage-dependent in these fungi. Interestingly, C-terminal or internal GFP fusions of VapA in *A. nidulans* resulted in non-functional or unstable proteins, in contrast to *S. pombe*, where internally tagged Scs2 remains functional [34]. Consistent with our findings, N-terminal GFP tagging has also been reported to produce functional VAP proteins in *S. cerevisiae* and *M. oryzae*.

Having established that VapA is a *bona fide* VAP homologue essential for proper growth, we next sought to understand the basis of the growth defect observed in the Δ*vapA* mutant. Since the genetic deletion of *vapA* had no apparent effect on unipolar germination or the initial stages of germling development, we hypothesized that the observed arrest in colony growth—including the failure to form mature mycelium and conidiospores—might stem from a defect in maintaining polarized growth. To test this, we examined the subcellular steady-state localization of apically localized protein cargoes involved in polarized growth, such as DnfA, DnfB, SynA, and ChsB. These proteins traffic to the hyphal tips via conventional, Golgi-dependent secretion, and they maintain their apical localization through constitutive endocytosis and recycling to the plasma membrane (PM). Our results revealed that, although secretion to the PM was not significantly impaired in the absence of VapA, apical cargoes related to lipid homeostasis (DnfA and DnfB, but also SynA) became depolarized, redistributing into the subapical segment of the PM, likely due to diffusion from the apical region. This cargo depolarization phenotype resembled what we had previously observed in mutants defective in apical endocytosis, such as Δ*ap2^σ^*or *thiAp-slaB* [36, 37]. Given that the AP-2 adaptor complex is a key regulator of cargo endocytosis at the hyphal tip, we hypothesized that VapA might be required for proper apical recruitment or activity of AP-2. Our data supported this idea: the absence of VapA phenocopied the effects of AP-2 dysfunction, affecting not only polarized growth and the distribution of ergosterol and phosphoinositides (PIs) in the PM, but also the apical localization of endocytic effectors such as SlaB and SagA. Thus, the simplest explanation for the cellular and developmental phenotypes observed in the Δ*vapA* mutant is that VapA, an ER-PM tethering protein, plays a critical role in facilitating AP-2-mediated endocytosis of lipid homeostasis related cargoes at the apical region. This function is likely essential for the polarized recycling of membrane proteins, and consequently, for maintaining long-term hyphal tip growth and colony development.

How could VapA affect AP-2 function? Although VapA and AP-2 do not colocalize (**Figure 4F**), suggesting no physical interaction, our data indicate that VapA may influence AP-2 function indirectly through its critical role in specific lipid partitioning at the PM, particularly at the apical region. Previous studies across various systems have established VapA as essential for proper lipid organization at the PM. Here we have provided compelling evidence supporting this role also in *A. nidulans*, including: a) altered filipin staining and increased sensitivity to itraconazole, indicating disrupted ergosterol partitioning at the PM, b) changes in the localization of pleckstrin homology (PH) domain markers, consistent with altered distribution of phospholipids such as PI4P in the Golgi and PI(4,5)P_2_ at the PM, c) mislocalization of the Sac1 phosphatase consistent with overaccumulation of PI4P and PI(4,5)P_2_, as all of the aforementioned changes are exhibited in the Δ*vapA* mutant.

Given the role of VapA in polarized cargo trafficking, we investigated whether it also affects the localization of non-polarized cargoes, such as the purine transporter UapA, the proton pump ATPase PmaA, and the alkaline pH sensor PalI. These proteins are not involved in apical cell expansion but are key for environmental sensing and adaptation, and PmaA is in fact necessary for growth via its pleiotropic action of pH homeostasis and nutrition. Notably, we previously showed that such non-polarized cargoes bypass the Golgi and likely reach the PM through alternative routes, possibly involving ER-PM contacts.

Importantly, in the present study, we provide direct evidence against this hypothesis. All three non-polarized cargoes tested trafficked normally to the PM in the Δ*vapA* mutant. Furthermore, toxic analogue growth assays indicated that other nutrient transporters remained functionally active in the absence of VapA. Supporting this, none of the ER-PM contact site mutants examined showed growth defects across a range of nutrients (**Figure 1C**), reinforcing that non-polarized nutrient transporters do not rely on ER-PM contacts for PM localization. In fact, we observed an increase in the steady-state PM levels of non-polarized cargoes in the Δ*vapA* mutant (see fluorescence intensity in **Figure 8B**). A similar increase in UapA translocation to the PM has also been reported in yeast Δtether strains [82]. These findings are consistent with recent work in *S. pombe* suggesting that ER-PM contacts can act as a physical barrier to vesicular secretion and protein exocytosis [34, 35].

Altogether, our results indicate that non-polarized trafficking to the PM in *A. nidulans* is independent of both the Golgi and ER-PM contact sites. VapA, although not physically associated with AP-2, may affect AP-2 function indirectly by shaping the lipid landscape of the PM. This altered lipid environment, especially perturbations in PI(4,5)P₂ distribution, could impair the formation or function of AP-2-dependent endocytic sites, thereby influencing the polarized trafficking of specific cargoes.

## MATERIALS AND METHODS

### Media, strains, growth conditions and transformation

Standard complete and minimal media for *A. nidulans* were used (FGSC, http://www.fgsc.net). Media and chemical reagents were obtained from Sigma-Aldrich (Life Science Chemilab SA, Hellas) or AppliChem (Bioline Scientific SA, Hellas). Glucose 1% (w/v) or fructose 0.1% (w/v) was used as carbon source. NH_4_^+^(di-ammonium tartrate, (NH4)_2_C_4_H_4_O) or NaNO_3_ were used as nitrogen sources at 10 mM. In growth tests with salt addition, KCl and NaH_2_PO_4_ were added at a final concentration of 0.6 M and 0.5 M respectively. Nucleobases and toxic analogs were used at the following final concentrations: 5-fluorouracil (5-FU), 8-azaguanine (AZG), 5-fluorocytosine (5-FC) and oxypurinol (OX) at 100 μM [84]; uric acid at 0.5mM. Itraconazole (ITZ) was used at a final concentration of 0.25 mg/L [11]. Thiamine hydrochloride was used at a final concentration of 10-20 μM as a repressor of the *thiA_p_* promoter in growth tests, microscopy or Western blot analysis [47]. *A. nidulans* transformation was performed by generating protoplasts from germinating conidiospores using TNO2A7 [85] or other *nkuA* DNA helicase deficient strains, that allow in-locus integrations of gene fusions via auxotrophic complementation. Integrations of gene fusions with fluorescent tags (GFP/mRFP/mCherry), promoter replacement fusions (*gpdA_p_/thiA_p_)* or deletion cassettes were selected using the *A. nidulans* marker para-aminobenzoic acid synthase (*pabaA*) or the *A. fumigatus* markers orotidine-5-phosphate-decarboxylase (AF*pyrG*, Afu2g0836), GTP-cyclohydrolase II (AF*riboB*, Afu1g13300) or a pyridoxine biosynthesis gene (AF*pyroA*, Afu5g08090), resulting in complementation of the relevant auxotrophies. GFP-tagged versions of VapA were firstly introduced in AN4406/*vapA* genomic locus of TNO2A7 strain, but failed to complement wild-type VapA. To identify a functional GFP-tagged VapA, plasmids overexpressing GFP-VapA, VapA-GFP-TM and VapA-ΔΤΜ-GFP under the *gpdA_p_* promoter, were introduced in ΔvapA*(AFpyrG)* pantoB100. GFP-VapA successfully complemented the null mutant (**Figure2B**). To that end, the plasmid carrying *gpdA_p_*-GFP-VapA was used for transformation of Δ7 (*ΔfurD::riboB ΔfurA::riboB ΔfcyB::argB ΔazgA ΔuapA ΔuapC::AfpyrG ΔcntA::riboB pabaA1 pantoB100*), based on complementation of the pantothenic acid auxotrophy pantoB100 [73]. Transformants were verified by PCR and growth test analysis. Combinations of mutations and fluorescent epitope-tagged strains were generated by standard genetic crossing and progeny analysis. *E. coli* strains used were DΗ5a. *A. nidulans* strains used are listed in **Supplementary Table S1-Strain List.** There is also a list containing annotations for genes referenced in the present work **Supplementary Table S2-Annotations**.

### Nucleic acid manipulations and plasmid constructions

Genomic DNA extraction was performed as described in FGSC (http://www.fgsc.net). Plasmid preparation and DNA gel extraction were performed using the Nucleospin Plasmid and the Nucleospin Extract II kits (Macherey-Nagel, Lab Supplies Scientific SA, Hellas). Restriction enzymes were from NEB (New England Biolabs, Bioline Scientific SA, Hellas). DNA sequences were determined by Eurofins-Genomics (Vienna, Austria). Conventional PCRs reactions were performed with KAPA Taq DNA polymerase (Kapa Biosystems, Lab Supplies Scientific). High-fidelity amplification of products and site-directed mutagenesis were performed with Kapa HiFi polymerase (Kapa Biosystems, Lab Supplies Scientific). Gene cassettes were generated by sequential cloning of the relevant fragments in the pGEM-T plasmid (Promega), which served as template to PCR-amplify the relevant linear cassettes. The genomic sequences for AN4406/*vapA,* AN9149/*tcbA*, AN5624/*tcbB*, AN2477/*istA*, AN7165/*istB* were retrieved from FungiDB (https://fungidb.org/fungidb/app). For AN4406/*vapA* deletion, several constructs were generated using pabaA, AF*pyroA* and AF*pyrG* genes. To construct VapA-GFP-TM and VapA-ΔΤΜ-GFP, SOSUI-transmembrane helix prediction tool (https://harrier.nagahama-i-bio.ac.jp/sosui/mobile/) was used, to identify a 22 amino acids transmembrane helix on the C-terminus of VapA (258-280aa, AGVPVRIVAGLCLLSFLIAYFFF). For the overexpression of GFP-tagged versions of *vapA,* a modified pGEM-T-easy vector was used, carrying a version of the gpdA promoter, the trpC 30 termination region, and the panB selection marker [86]. Oligonucleotides used for cloning are listed in **Supplementary Table S3-Primer List.**

### Phylogenetic tree

Protein sequences were retrieved from the UniProt database (https://www.uniprot.org/) for a representative selection of model organisms. All sequences were manually curated to ensure orthology and high annotation quality. Multiple sequence alignment was conducted using the MAFFT online server [87] at https://mafft.cbrc.jp/alignment/server/, applying the E-INS-i algorithm. Alignment settings included two iterative refinement cycles, and the “Try to align gappy regions anyway” option was selected to preserve informative gaps. The final alignment comprised 23 sequences with 557 amino acid positions. Phylogenetic tree was constructed using the IQ-TREE web server v1.6.12 [88, 89] with integrated ModelFinder [90] for automated substitution model selection. The best-fitting model according to the Bayesian Information Criterion (BIC) was LG+I+G4, which includes a general amino acid replacement matrix (LG), a proportion of invariable sites (I), and gamma-distributed rate heterogeneity across four discrete categories (G4). Maximum likelihood (ML) tree inference was performed under this model, and ultrafast bootstrap approximation (UFBoot2) was applied with 1,000 replicates to assess branch support [91]. The resulting unrooted ML tree was rooted using the plant clade as the outgroup. The tree was visualized and annotated using FigTree v1.4.4 (http://tree.bio.ed.ac.uk/software/figtree/). **Supplementary Table S4** compiles the uniport IDs used to generate the tree. Note that for AN4406/VapA, the sequence was retrieved from FungiDB (https://fungidb.org/fungidb/app), as there were missing residues on the C-tail at the Uniprot sequence (Q5B4X4).

### Protein extraction and Western blots

Protein extraction and western blotting Total protein extraction was performed as previously described by [69, 70]. For strains shown in **Supplementary Figure S1 and Supplementary Figure S3**, dry mycelia from cultures grown in minimal liquid cultures supplemented with 10 mM NH_4_^+^, and the required auxotrophies were used (culture conditions: 25°C for 16 h,). For *thiA_p_*-*tcbB,* thiamine hydrochloride was added at a final concentration of 20 μM. Total proteins (50 μg, estimated by Bradford assays) were separated in a 10 % (w/v) poly acrylamide gel and then transferred on PVDF membranes (GE Health care Life Sciences, Amersham). Immunodetection was performed with an anti-FLAG antibody (Agrisera, AAS15 3037) for *thiA_p_*-FLAG-*tcbB* or an anti-GFP antibody (11814460001, Roche Diagnostics) for *gpdA_p_*-GFP-VapA constructs. Then an HRP-linked anti body (7076, Cell Signaling Technology Inc.) was used in both cases. Colloidal Coomassie G-250 Staining for Proteins (CBB) was employed as described in Dyballa and Metzger, 2009, to ensure equal loading. For *thiA_p_*-FLAG-*tcbB* Western blot, an anti-actin monoclonal (C4) antibody (SKU0869100-CF, MP Biomedicals, Europe), was also employed. Blots were developed using the Lumi Sensor Chemiluminescent HRP Substrate kit (Genscript, United States) and SuperRX Fuji medical X-Ray films (Fuji FILM, Europe).

### Fluorescence microscopy

Conidiospores were incubated overnight in glass bottom 35mm μ-dishes (ibidi, Lab Supplies Scientific SA, Hellas) in liquid minimal media containing 1% (w/v) glucose, supplemented with NH_4_^+^, and the required auxotrophies, for 16–22h at 25°C. For following the subcellular trafficking and localization of DnfA, DnfB, ChsB, UapA, PmaA, PalI, these cargoes were expressed under their native promoter. The *uapA* promoter can be tightly repressed in the presence of 10 mM ammonium tartrate supplied as a nitrogen source in the growth medium, and derepressed in the presence of nitrate. mCherry-SynA on wt and Δ*ap2σ* strain was expressed under its native promoter. GFP-SynA was expressed under the regulatable *alcA* promoter in wt and Δ*vapA* strains. *AlcA_p_* is repressed on 1% w/v glucose and derepressed upon shift to 0.1% w/v fructose for 4-5h. For following the subcellular localization of de novo GFP-OshB, the transcription was induced upon shift to 0.1% w/v fructose and 0.4% v/v ethanol containing media for 4 h. Selected proteins (*thiA_p_-myoA, thiA_p_-slaB*) expressed under the *thiA* promoter, were transcriptionally repressed in the presence of 10μM thiamine in the growth media. SynaptoRed C2 (Equivalent to FM4-64) (Biotium) staining took 5-8 min on ice at a final concentration of 4μM, and observed at 5min and 25min post-washes. CellTracker™ Blue

CMAC (InvitrogenTM) was used according to [44]. FilipinIII (Cayman Chemical) staining took place 5min, room temperature at 1 μg/ml final concentration [36]. Calcofluor white was used at 0,001% w/v final concentration, for 5min, RT [36]. Images from widefield microscopy were obtained using an inverted Zeiss Axio Observer Z1 equipped with the white light pE-400 Illumination System (https://www.coolled.com/products/pe-400) and a Hamamatsu ORCA-Flash 4 camera. All widefield z-stack images were deconvolved with Huygens Essential version 23.10 (Scientific Volume Imaging, The Netherlands, http://svi.nl). Technical replicates correspond to different hyphal cells observed within each sample, while biological replicates refer to different samples. Each experiment has been conducted at least two times.

### Image processing and Statistical analysis

Images of fungal colonies from growth assays were processed for comparative presentation using Adobe Photoshop CS4 Extended (version 11.0.2). Individual colonies representing different *Aspergillus nidulans* strains were isolated from Petri dishes photographs, using the “layer via Copy” tool, allowing rearrangement of colonies from different plates into a single composite figure. (Figure 1C, 2A, 3A, 5A, 6C, 9A, Supplementary Figure S1, Supplementary Figure S2). Adjustments to brightness and contrast were applied to improve visibility across samples. All image processing steps were limited to figure preparation and did not affect the experimental analysis. Each growth test has been conducted at least two times. Microscopy images were processed and analyzed using Fiji/ImageJ [92]. Maximum intensity projections, contrast adjustments, region-of-interest (ROI) selection, color channel merging, and scale bar insertion were performed to enhance image clarity for figure presentation. These manipulations were applied uniformly, after any fluorescence intensity measurement and did not alter the underlying fluorescence signals used for quantification. Additional image annotation and formatting were performed using Adobe Photoshop CS4 Extended (version 11.0.2). To quantify cortical ER fluorescence labeled with Sec63-mCherry (**Figure 4A**), the polygon selection tool in Fiji was used to define a ROI approximately 10–12 μm from the hyphal tip (ROI_total). A second, smaller ROI within this region was drawn to measure the cytoplasmic signal (ROI_intracellular). Fluorescence intensities were normalized to area, and cortical ER fluorescence per μm was calculated by subtracting the intracellular signal from the total: Cortical ER Fluorescence/Area = (Total IntDen / Area) – (Intracellular IntDen / Area). The resulting values were compared between wild-type (WT) and Δ*vapA* strains using an unpaired t-test. The mean ± SEM for WT was 90.63 ± 12.64 (N = 13), and for ΔvapA was 107.7 ± 15.29 (N = 17), with t = 0.8263, degrees of freedom (df) = 28, and p = 0.4156. To assess ER–plasma membrane contact sites marked by TcbA-GFP (**Figure 4A**), a segmented line approximately 0–12 μm in length was drawn along the plasma membrane at the hyphal tip. Fluorescence intensity profiles were generated using Fiji/ImageJ, and regions with intensity values greater than 1000 units were considered TcbA-enriched. The length of TcbA-positive membrane was divided by the total measured membrane length to yield a ratio per cell. WT cells showed a mean ratio of 0.6575 ± 0.07126 (N = 17), and ΔvapA cells showed 0.5415 ± 0.06840 (N = 20), with t = 1.171, df = 35, and p = 0.2495. For SedV and PH^OSBP^ (**Figure 4A**), fluorescence intensity was measured using a polygon ROI starting approximately 8 μm from the hyphal tip, with each value corresponding to a single hypha (technical replicate). For SedV, WT had a mean intensity of 875.4 ± 65.84 (N = 24) and Δ*vapA* had 706.7 ± 47.21 (N = 28), yielding t = 2.123, df = 50, and p = 0.0387. For PH^OSBP^, Welch’s t-test was used due to unequal variances: WT had a mean of 1835 ± 409.7 (N = 20) and Δ*vapA* had 4207 ± 840.0 (N = 20), with t = 2.538, df = 27, and p = 0.0172. All statistical analyses for the Sec63, TcbA, SedV, and PH^OSBP^ experiments were performed using GraphPad Prism3 (GraphPad Software, San Diego, CA, USA). Full datasets and statistical outputs for these comparisons are available in Data_Fig4. For quantification of apical fluorescence gradients (**Figures 5C, 7 and Supplementary figure S4**), a line scan analysis was performed. A straight line was drawn from the hyphal tip extending 5 μm into the cell interior. Fluorescence intensity along this line was measured using the line profile tool in Fiji. This procedure was repeated for each hypha, and the average intensity at each position along the line was calculated for each strain. For region-specific analysis, intensity values were binned into 1 μm intervals (i.e., 0–1 μm, 1–2 μm, 2–3 μm, 3–4 μm, and 4–5 μm). Within each bin, fluorescence values for each strain were pooled, and pairwise comparisons were conducted between WT and Δ*vapA*, as well as WT and Δ*ap2σ*. Statistical outputs and source data are included in Data_Fig5 and Data_Fig7. A similar method was applied in **Figures 4B, 4C, 5E, 6A, and 7** to quantify cortical fluorescence intensity surrounding the plasma membrane at the hyphal tip. In these experiments, segmented lines were drawn along the plasma membrane covering distances of 0–10 μm or 0–12 μm, depending on the figure. Fluorescence values along these lines were binned by 1 μm and analyzed similarly to the apical gradient measurements. Datasets and corresponding statistical analyses for these figures are provided in Data_Fig4, Data_Fig5, Data_Fig6, and Data_Fig7. For FM4-64 internalization experiments (Figure 6D), the polygon selection tool in Fiji was used to define a ROI approximately 10 μm from the hyphal tip (ROI_total). A second, smaller ROI within this region was drawn to measure the cytoplasmic signal (ROI_intracellular). Fluorescence intensities were normalized to area, and PM fluorescence per μm was calculated by subtracting the intracellular signal from the total: PM Fluorescence/Area = (Total IntDen / Area) – (Intracellular IntDen / Area). For our analysis the Intracellular IntDen/Area corresponds to the Cytosolic fluorescence per μm. The resulting values were grouped for the two conditions used: 5 and 25 min FM4-64 internalization respectively. Unpaired t-test to compare the PM and cytosolic fluorescence intensity of wild-type (WT) and Δ*vapA* was employed. Notably, only the comparison of the cytosolic fluorescence intensity at 25 min, proved to be statistically significant (p<0.05, p= 0.0308, t=2.201 df=75, N_WT_= 38, N_ΔvapA_=39). Statistical analyses for the apical fluorescence gradient and cortical fluorescence intensity measurement, were performed using a custom Python script (Python v3.10) utilizing pandas, NumPy, and SciPy.Stats libraries [93–95]. A 95% confidence interval was applied for all tests. Prior to hypothesis testing, data normality was assessed using the Shapiro–Wilk test, and homogeneity of variances was evaluated using Levene’s test. If both assumptions were met, an unpaired t-test was used; if variances were unequal, Welch’s t-test was applied; and if normality was violated in either group, the non-parametric Mann–Whitney U test was employed. All fluorescence intensity plots were generated using GraphPad Prism3. For co-localization analysis, Pearson’s correlation coefficient (PCC) was calculated using the BIOP JACoP plugin in Fiji [96]. PCC values were computed from maximum intensity projections of deconvolved z-stacks, using manually defined ROIs that encompassed each hypha. Statistical analysis and visualization of PCC values were performed using GraphPad Prism, and a one-sample t-test with a 95% confidence interval was used to determine significance.

## Data Availability Statement

Strains and plasmids are available upon request. The authors affirm that all data necessary for confirming the conclusions of the article are present within the article, figures, and tables.

## ACKNOWLEDGMENTS

This work was supported by an H.F.R.I. research grant (KE18458). Xenia Georgiou was supported by *Fondation Santé.* We thank Dr. Ioannis Adamakis for providing us with Calcofluor white dye.

## SUPPLEMENTAL MATERIAL

All supplemental data for this article are available online at www.microbialcell.com.

## Abbreviations

PM: Plasma membrane
ER: Endoplasmic reticulum
wt: wild type
cER: cortical ER
TM: transmembrane
PI(4,5)P_2_: Phosphatidylinositol (4,5)-bisphosphate
PI4P: Phosphatidylinositol 4-phosphate

## Supplementary information

### Supplementary Figures

**Supplementary figure S1:**
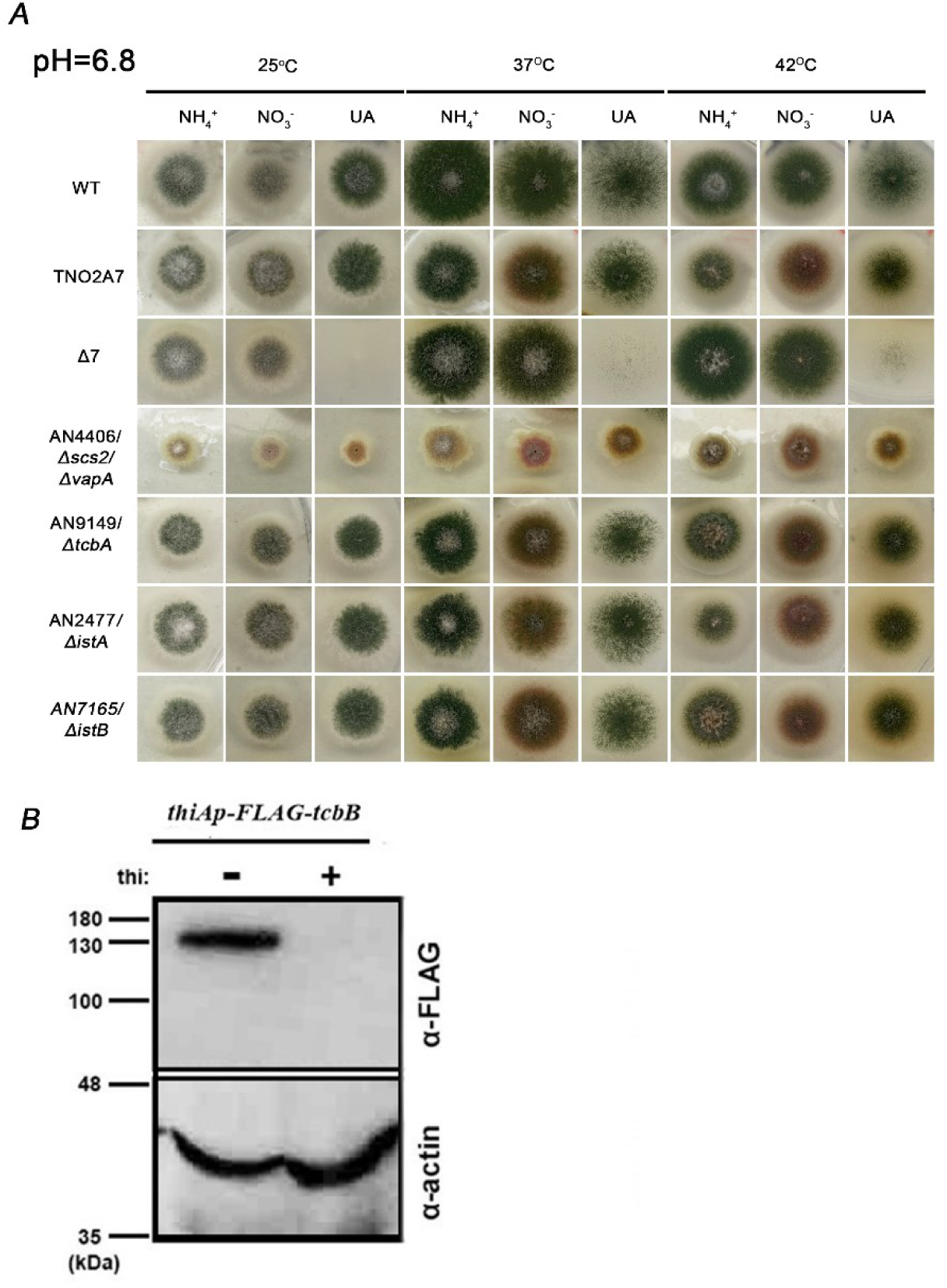
Extended growth test, part of which is presented in **Figure1C**. Strains carrying single total deletions of genes encoding VapA/VAP, tricalbins or Ist2 proteins (Δ*vapA*, Δ*tcbA*, Δ*istA* and Δ*istB*). Growth of doubly knockout mutant *ΔistA/ΔistB* is also shown. A standard isogenic wild-type strain (wt) and a strain carrying multiple deletions in genes encoding purine/pyrimidine-related transporters are used as controls (Δ*furD* Δ*furA* Δ*fcyB* Δ*uapA* Δ*uapC* Δ*azgA* Δ*cntA*, named Δ7). Strains are grown on minimal media with selected nitrogen sources (ammonium/NH_4_^+^ or nitrate/NO_3_^-^ or uric acid/UA), at pH=6.8, at 25 °C, 37°C, 42 °C. **B.** Western blot analysis of TcbB (AN5624) using the anti-FLAG antibody. In the absence of thiamine (−thi) from the growth medium, TcbB is expressed, while upon addition of thiamine (+thi) at the onset of conidiospore germination (ab initio repression), the expression of these proteins is tightly repressed. Equal loading and protein steady-state levels are normalized against the amount of actin, detected with an anti-actin antibody.

**Supplementary figure S2:**
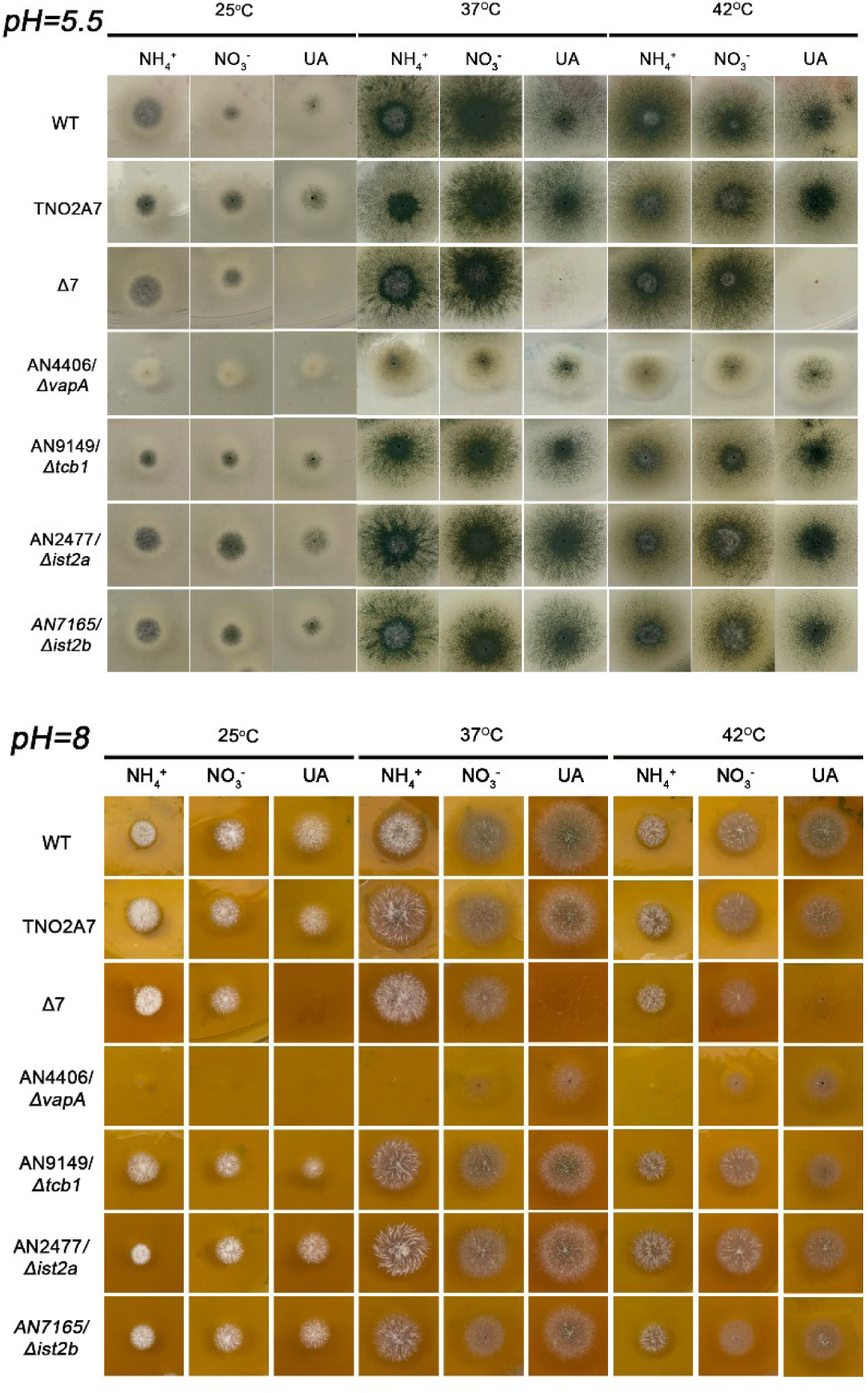
Growth phenotypes of strains carrying single total deletions of genes encoding VapA/VAP, tricalbins or Ist2 proteins (Δ*vapA*, Δ*tcbA*, Δ*istA* and Δ*istB*). Growth of doubly knockout mutant *ΔistA/ΔistB* is also shown. A standard isogenic wild-type strain (wt) and a strain carrying multiple deletions in genes encoding purine/pyrimidine-related transporters are used as controls (Δ*furD* Δ*furA* Δ*fcyB* Δ*uapA* Δ*uapC* Δ*azgA* Δ*cntA*, named Δ7). Strains are grown on minimal media with selected nitrogen sources (ammonium/NH_4_^+^ or nitrate/NO_3_^-^ or uric acid/UA), at pH=5.5 and 8, at 25 °C, 37°C, 42 °C.

**Supplementary figure S3:**
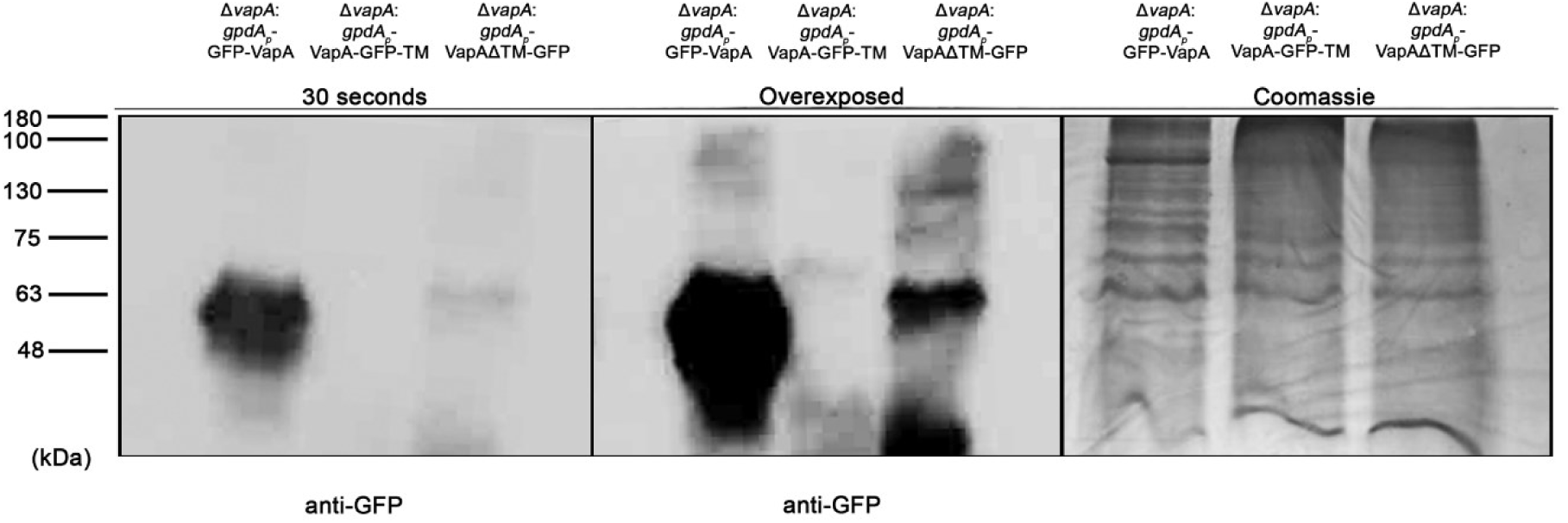
Western blot analysis of GFP-VapA, VapA-GFP-TM, VapA-ΔΤΜ-GFP, under the *gpdA* promoter in Δ*vapA* background using the anti-GFP antibody. Notice there is high expression of the VapA-GFP construct, estimated at 63 kDa, while VapA-GFP-TM is barely detected and VapA-ΔΤΜ-GFP exhibits increased degradation. Equal loading was achieved using Coomassie staining.

**Supplementary figure S4:**
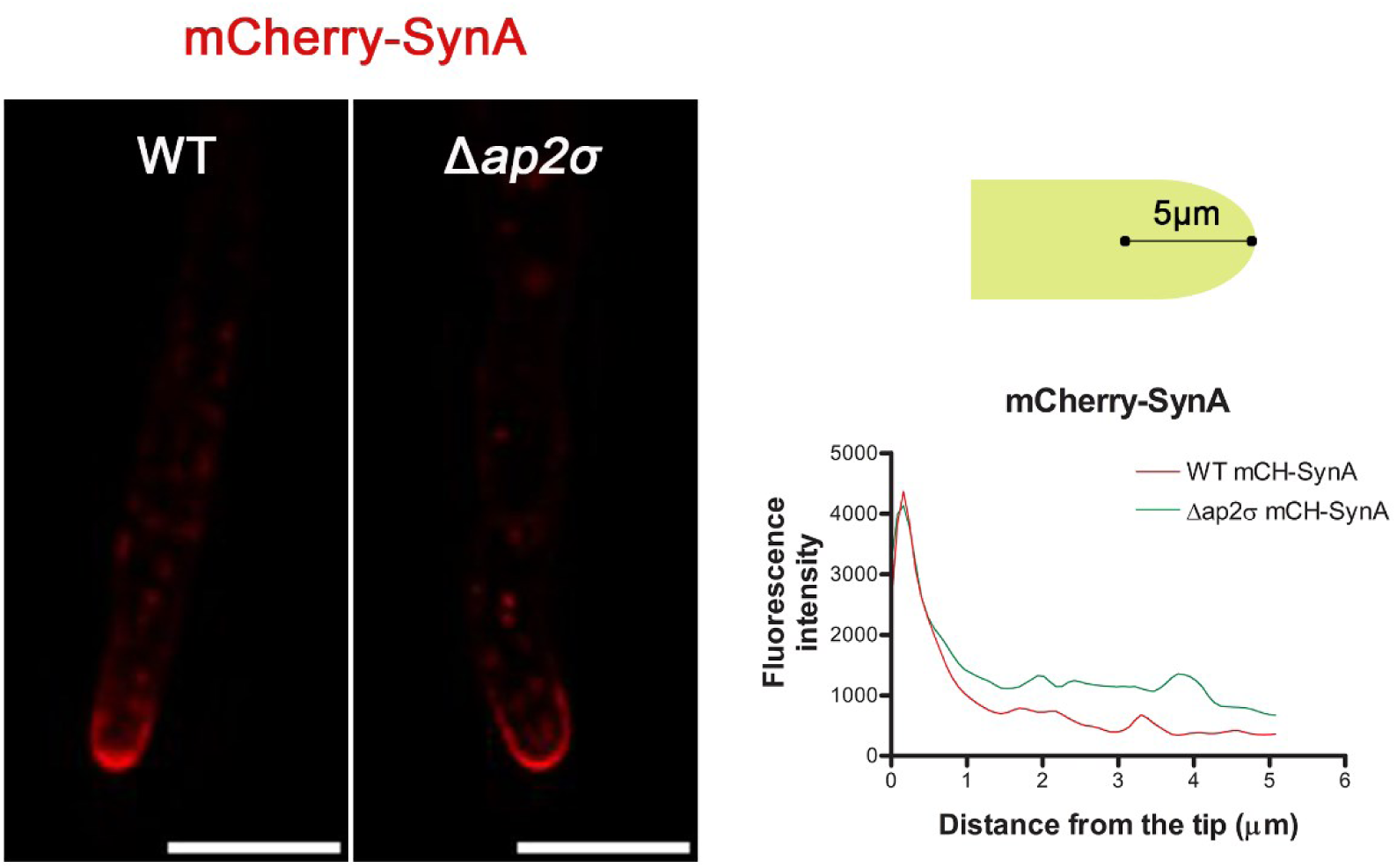
mCherry-SynA localization in wt and Δ*ap2^σ^*. Maximal intensity projections of deconvolved snap shots showing the localization of mCH-SynA in wt and Δ*ap2^σ^* background. Notice that SynA is unaffected on Δ*ap2^σ^.* Line plot showing SynA fluorescence intensity along the hyphal tip in wild-type (WT, red) and Δ*ap2^σ^*(green) strains. The x-axis represents the distance from the hyphal tip (μm), and the y-axis represents fluorescence intensity.

**Supplementary figure S5.**
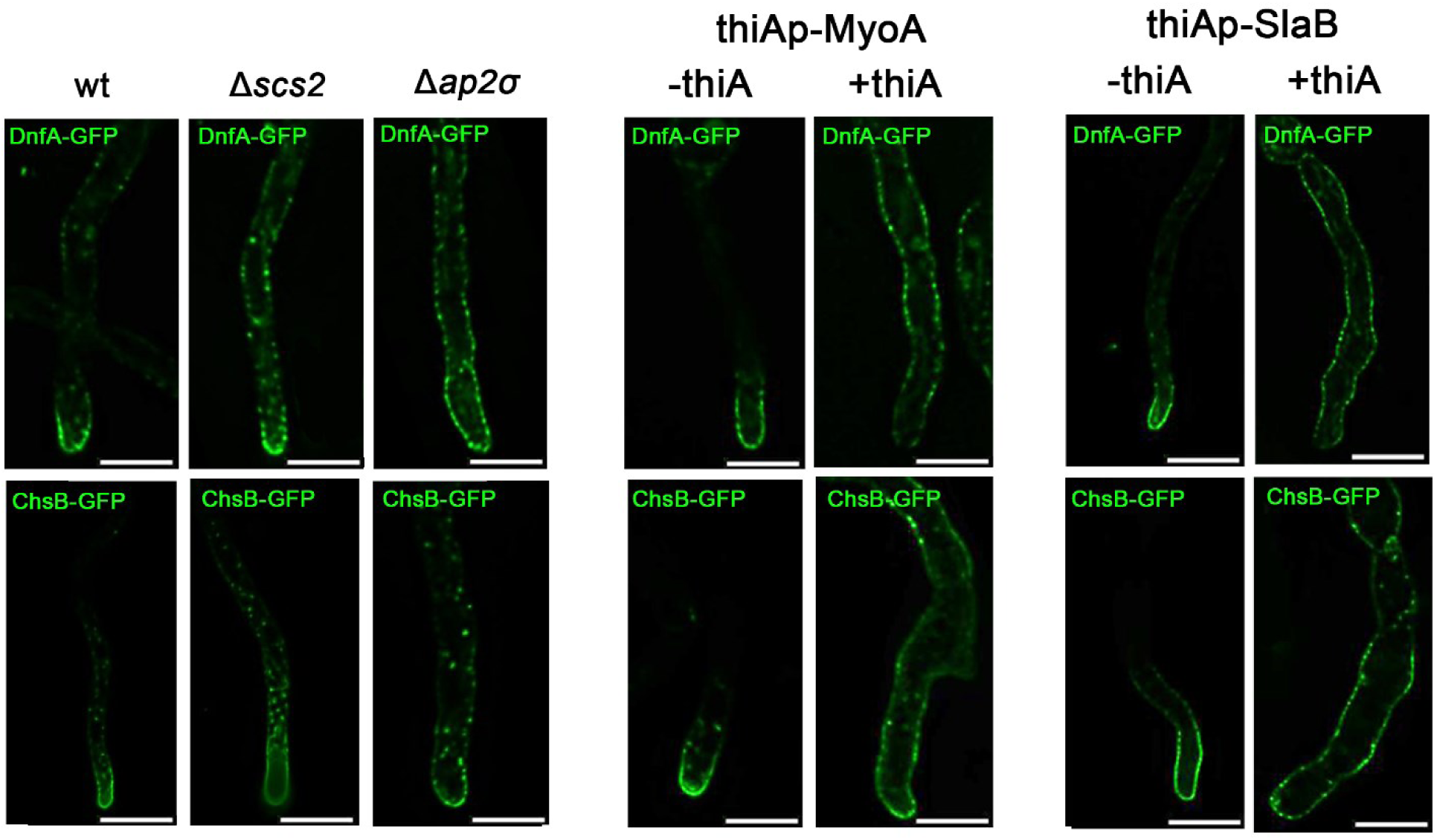
Comparison of DnfA and ChsB localization in Δ*vapA*, Δ*ap2^σ^, thiAp-myoA* and *thiAp-slaB*. Notice that upon repression of endocytosis factors, MyoA and SlaB, DnfA and ChsB have totally lost their apical localization and mark homogeneously the plasma membrane. In Δ*ap2^σ^*, only DnfA exhibits severe endocytosis defect, while ChsB remains apical yet labels intracellular loci. A similar phenotype for ChsB, was observed also in Δ*vapA*. Scale bars: 5 μm

### Supplementary Tables

**Supplementary Table S1:** Strain list.

| **Name** | **Genotype** | **Reference** |
| --- | --- | --- |
| **TNO2A7** | ΔnkuA::argB pyrG89 pyroA4 riboB2 | Nayak et al., 2006 |
| **wt** | pabaA1 | wild-type reference strain |
| **ΔvapA** | (AN4406)vapAΔ::AFpyrG nkuAΔ::argB pyroA4 riboB2 pyrG89 | This study |
| **ΔvapA** | (AN4406)vapAΔ::AFpyrG pantoB100 | This study |
| **ΔtcbA** | (AN9149)tcbAΔ::AFpyrG89 pyrG89 riboB2 pyroA4 nkuAΔ::argB | This study |
| **ΔistA** | (AN2477)ist2AΔ::AFriboB2 nkuAΔ::argB pyroA4 riboB2 pyrG89 | This study |
| **ΔistB** | (AN7165)ist2BΔ::AFpyrG nkuAΔ::argB pyroA4 riboB2 pyrG89 | This study |
| **thiAp-tcbB** | thiAp-FLAG-AN5624(TcbB)::AFriboB pyrG89 riboB2 pyroA4 nkuAΔ::argB | This study |
| **ΔistA/ΔistB** | (AN2477)ist2AΔ::AFriboB2 (AN7165)ist2BΔ::AFpyrG nkuAΔ::argB pyroA4 riboB2 pyrG89 | This study |
| **ΔtcbA/thiAp-tcbB** | AN9149(TcbA)$\Delta$::AFpyrG89 thiAp-FLAG-AN5624(TcbB)::AFriboB pyrG89 riboB2 pyroA4 nkuAΔ::argB | This study |
| **gfp-vapA** | GFP-vapA::AFpyrG nkuAΔ::argB pyroA4 riboB2 pyrG89 | This study |
| **vapA-ΔTM-gfp** | vapA-ΔTM-GFP::AFpyrG nkuAΔ::argB pyroA4 riboB2 pyrG90 | This study |
| **vapA-gfp-TM** | vapA-GFP-TM::AFpyrG nkuAΔ::argB pyroA4 riboB2 pyrG91 | This study |
| **vapA-gfp** | vapA-GFP::AFpyrG nkuAΔ::argB pyroA4 riboB2 pyrG92 | This study |
| **ΔvapA gpdA-gfp-vapA** | (AN4406)vapAΔ::AFpyrG pantoB100 pGEM-gpdA-GFP-(GA)x5-AN4406/vapA-trpC-panB | This study |
| **ΔvapA gpdA-vapA-ΔTM-gfp** | (AN4406)vapAΔ::AFpyrG pantoB100 pGEM-gpdA-AN4406/vapAΔTM-GFP-TM-trpC-panB | This study |
| **ΔvapA gpdA-vapA-gfp-TM** | (AN4406)vapAΔ::AFpyrG pantoB100 pGEM-gpdA-AN4406/vapAΔTM-GFP-trpC-panB | This study |
| **wt gpdA-gfp-vapA** | ΔuapA UapC::pyrG ΔazgA ΔFcyB::argB ΔFurD::riboB ΔFurA::riboB ΔcntA pabaA1 pantoB100 pGEM-gpdA-GFP-(GA)x5-AN4406/vapA-trpC-panB | This study |
| **wt mCherry-vapA** | pyrG89 argB2 pabaA1 pantoB100 pGEM gpdAp-mCherry-(GA)x2-AN4406/vapA-trpC-panB | This study |
| **ΔvapA gpdA-mCherry-vapA** | (AN4406)vapAΔ::AFpyrG pantoB100 gpdAp-mCherry-(GA)x2-AN4406/vapA-trpC-panB | This study |
| **wt gpdA-gfp-vapA sec63-mCherry** | sec63-mCherry::pyrG pGEM-gpdA-GFP-(GA)x5-AN4406/vapA-trpC-panB pabaA1 | This study |
| **wt gpdA-gfp-vapA mrfp-PHosbp** | pyroA4::[pyroA-gpdAmini-mrfp-PHOSBP] pGEM-gpdA-GFP-(GA)x5-AN4406/vapA-trpC-panB inoB2 wA4 | This study |
| **wt gpdA-gfp-vapA sec63-mCherry** | pyroA4::[pyroAmut::gpdAmini::mcherry::sedV] pGEM-gpdA-GFP-(GA)x5-AN4406/vapA-trpC-panB inoB2 wA4 | This study |
| **wt sec63-mCherry** | sec63-mCherry::pyrG pabaA1 pantoB100 | Buhler et al. |
| ***ΔvapA sec63-mCherry uapA-gfp*** | (AN4406)scs2Δ::AFpyrG sec63-mCherry::pyrG uapAΔ::uapA-GFPL::AFriboB2 pabaA1 pantoB100 | This study |
| ***wt tcbA-gfp*** | AN9149/(TcbA)-gfp::AFpyrG pyrG89 riboB2 pyroA4 nkuAΔ::argB | This study |
| ***ΔvapA tcbA-gfp*** | AN4406:: AFriboB AN9149 (TcbA)-gfp::AFpyrG pyrG89 riboB2 pyroA4 nkuAΔ::argB | This study |
| ***wt mrfp-PHosbp*** | MAD2013: [pyroA-gpdAmini-mrfp-PHOSBP]pyroA4 inoB2 niiA4 wA4 | Pantazopoulou et al., 2009 |
| ***ΔvapA dnfA-gfp mrfp-PHosbp*** | (AN4406)vapAΔ::AFpyrG dnfA-GFP::AFpyrG [pyroA-gpdAmini-mrfp-PHOSBP]pyroA4 pyroA4 pyrG89 wA4 | This study |
| ***wt mCherry-sedV*** | MAD2808: pyroA4::[pyroAmut::gpdAmini::mcherry::sedV] nkuAΔ::bar, wA4, niiA4 inoB2 | Pantazopoulou et al., 2009 |
| ***ΔvapA mCherry-sedV*** | (AN4406)vapAΔ::AFpyrG pyroA4::[pyroAmut::gpdAmini::mcherry::sedV] pantoB100 inoB2 | This study |
| ***wt AKL-mRFP*** | pGEM-gpdAp-mRFP<sub>AKL</sub>-trpC-panB pantoB100 biA1 | Galanopoulou et al., 2014 |
| ***ΔvapA AKL-mRFP*** | (AN4406)vapAΔ::AFpyrG pGEM-gpdAp-mRFP<sub>AKL</sub>-trpC-panB pyroA4 biA1 | This study |
| ***wt PHplcδ-gfp*** | [gpdA::gfp::PHdomPLCb12]-argBmut::argB2 pabaA1 yA2 | Pantazopoulou et al., 2009 |
| ***ΔvapA PHplcδ-gfp*** | (AN4406)vapAΔ::pabaA1 [gpdA::gfp::PHdomPLCb12]-argBmut::argB2 pabaA1 yA2 | This study |
| ***ΔvapA H1-mRFP*** | (AN4406)vapAΔ::AFpyrG LO:1516 H1-mRFP::AFriboB pyroA4 pyrG89 riboB2 nkuAΔ::argB | This study |
| ***H1-mRFP*** | LO:1516 H1-mRFP::AFriboB nkuAΔ::argB pyrG89 pyroA4 riboB2 | Ramón et al., 2000 |
| ***wt dnfA-gfp*** | dnfA-GFP::AFpyrG nkuAΔ::argB pyrG89 pabaA1 pyroA4 | Schultzhaus et al., 2015 |
| ***ΔvapA dnfA-gfp*** | (AN4406)vapAΔ::AFpyrG dnfA-GFP::AFpyrG pyroA4 | This study |
| ***wt dnfB-gfp*** | dnfB-GFP::AFpyrG nkuAΔ::argB pyrG89 pabaA1 pyroA4 | Schultzhaus et al., 2015 |
| ***ΔvapA dnfB-gfp*** | (AN4406)vapAΔ::AFpyrG dnfB-GFP::AFpyrG nkuAΔ::argB pyrG89 pabaA1 pyroA4 | This study |
| ***ΔdnfA*** | dnfAΔ::AFriboB nkuAΔ::argB pyrG89 pyroA4 | Schultzhaus et al., 2015 |
| ***ΔdnfB*** | dnfBΔ::AFriboB nkuAΔ::argB pyrG89 pyroA4 | Schultzhaus et al., 2015 |
| ***Δap2σ*** | Ap2σΔ::AFriboB nkuAΔ::argB pyroA4 riboB2 pyrG89 | Martzoukou et al., 2017 |
| ***Δap2σ/ΔvapA dnfA-gfp*** | (AN0722)Ap2σΔ::AFriboB (AN4406)vapAΔ::AFpyrG dnfA-GFP::AFpyrG pyroA4 | This study |
| ***wt alcAp-synA-gfp*** | alcAp-GFP-synA::AFpyrG nkuAΔ::argB pyroA4 riboB2 pyrG89 | Dimou et al., 2020 |
| ***ΔvapA alcAp-synA-gfp sec63-mCherry*** | (AN4406)vapAΔ::AFpyroA alcAp-GFP-synA::AFpyrG sec63-mCherry::pyrG pyroA4 riboB2 pyrG89 | This study |
| ***wt gfp-chsB*** | GFP-chsB::AFpyrG pyrG89 pabaA1 | Fukuda et al., 2009 |
| ***ΔvapA gfp-chsB*** | (AN4406)vapAΔ::AFpyrG GFP-chsB::AFpyrG | This study |
| ***wt mCherry-synA*** | fwA1 pyrG89 pyroA4 nicA2 nkuAΔ::argB AFpyG-mcherry-synA yA::AFpyroA tpmAp-gfp-tpmA | Martzoukou et al., 2017 |
| ***Δap2σ mCherry-synA*** | ap2Δ mCherry-SynA GFP-TmpA yA2 (2) | Martzoukou et al., 2017 |
| ***Δap2σ dnfA-gfp*** | dnfA-GFP::AFpyrG Ap2σΔ::AFriboB nkuAΔ::argB pyrG89 pyroA4 (7) | Martzoukou et al., 2017 |
| ***Δap2σ dnfB-gfp*** | ap2σΔ::AFriboB DnfB-GFP::AFpyrG nkuAΔ::argB pyrG89 pyroA4 (2) | Martzoukou et al., 2017 |
| ***Δap2σ gfp-chsB*** | GFP-chsB::AFpyrG ap2σΔ::AFriboB nkuAΔ::argB pyrG89 riboB2 pyroA4 | Martzoukou et al., 2017 |
| ***wt ap2σ-mrfp*** | ap2σ-(5xGA)mRFP::AFpyrG nkuAΔ::argB pyroA4 riboB2 pyrG89 | This study |
| ***ΔvapA ap2σ-mrfp*** | (AN4406)vapAΔ::AFpyrG ap2σ-(5xGA)mRFP::AFpyrG pyroA4 | This study |
| ***ΔvapA/Δerg11A/thiAp-erg11B*** | ΔAN4406::AFriboB AN1901Δ::AFpyrG thiAp-AN8283::AFriboB nkuAΔ::argB pyrG89 pyroA4 riboB2 pantoB | This study |
| ***ΔvapA/Δerg4A/Δerg4B*** | ΔAN4406::AFriboB AN2684Δ::AFpyrG AN10648Δ::AFriboB nkuAΔ::argB pyrG89 pyroA4 riboB2 | This study |
| ***ΔvapA/thiAp-pisA*** | ΔAN4406::AFpyrG thiAp-AN0913::AFpyrG nkuAΔ::argB pyrG89 pyroA4 riboB2 pantoB | This study |
| ***ΔvapA/thiAp-basA*** | ΔAN4406::AFpyrG thiAp-AN0640::AFpyrG nkuAΔ::argB pyrG89 pyroA4 riboB2 pantoB | This study |
| ***Δerg11A/thiAp-erg11B*** | AN1901Δ::AFpyrG thiAp-AN8283::AFriboB nkuAΔ::argB pyrG89 pyroA4 riboB2 | Dionysopoulou et al., 2021 |
| ***Δerg4A/Δerg4B*** | AN2684Δ::AFpyrG AN10648Δ::AFriboB nkuAΔ::argB pyrG89 pyroA4 riboB2 | Dionysopoulou et al., 2021 |
| ***thiAp-pisA*** | thiAp-AN0913::AFpyrG nkuAΔ::argB pyrG89 pyroA4 riboB2 | Dionysopoulou et al., 2021 |
| ***thiAp-basA*** | thiAp-AN0640::AFpyrG nkuAΔ::argB pyrG89 pyroA4 riboB2 | Dionysopoulou et al., 2021 |
| ***ΔvapA myoA-gfp*** | ΔAN4406::AFpyrG AN1558(myoA)-(5xGA)GFP::AFpyrG nkuAΔ::argB pyrG89 pyroA4 riboB2 pantoB | This study |
| ***wt myoA-gfp*** | AN1558(myoA)-(5xGA)GFP::AFpyrG nkuAΔ::argB pyrG89 pyroA4 riboB2 | Martzoukou et al., 2017 |
| ***ΔvapA slab-gfp*** | ΔAN4406::AFpyrG slaB-GFP::AFpyrG pyrG pyroA4 riboB2 argB2 Δnku::argB | This study |
| ***wt slab-gfp*** | slab-GFP::AFpyrG pyrG pyroA4 argB2, Δnku::argB | Araujo-Bazán et al., 2008 |
| ***ΔvapA abpA-dsRed*** | ΔAN4406::AFpyrG [abpA-mRFP::AFpyrG] pabaA1 | This study |
| ***wt abpA-dsRed*** | yA2, pabaA1, pyrG89, [abpA-mRFP::AFpyrG] | Araujo-Bazán et al., 2008 |
| ***ΔvapA saga-gfp*** | ΔAN4406::AFpyrG sagA/end3-GFP::AFpyrG nkuAΔ::argB riboB2 pyroA4 pyrG89 pantoB | This study |
| ***wt sagA-gfp*** | sagA/end3-GFP::AFpyrG nkuAΔ::argB riboB2 pyroA4 pyrG89 | Karachaliou et al., 2013 |
| ***Δap2σ abpA-dsRed*** | (AN0722)Ap2σ $\Delta$::AFriboB [abpA-mRFP::AFpyrG] pabaA1 pyrG89 | Martzoukou et al., 2017 |
| ***Δap2σ slab-gfp*** | ap2σ $\Delta$::AFriboB slab-GFP::AFpyrG nkuAΔ::argB pyroA4 | Martzoukou et al., 2017 |
| ***Δap2σ saga-gfp*** | ap2σ $\Delta$::AFriboB sagA-GFP::AFpyrG pabaA1 pyroA4 | Martzoukou et al., 2017 |
| ***Δap2σ myoA-gfp*** | ΔAN4406::AFriboB AN1558(myoA)-(5xGA)GFP::AFpyrG nkuAΔ::argB pyrG89 pyroA4 riboB2 | Martzoukou et al., 2017 |
| ***thiAp-myoA dnfA-gfp*** | DnfAGFP::AFpyrG thiAp-AN1558(myoA)::AFpyrG nkuAΔ::argB pyrG89 pyroA4 riboB2 | Martzoukou et al., 2017 |
| ***thiAp-slab dnfA-gfp*** | dnfA-GFP::AFpyrG slab::thiAp-slab::AFriboB nkuAΔ::argB pyroA4 riboB2 pyrG89 | Martzoukou et al., 2018 |
| ***thiAp-myoA gfp-chsB*** | thiAp-myoA::AFpyrG GFP-chsB::AFpyrG pabaA1 | Martzoukou et al., 2018 |
| ***thiAp-slab gfp-chsB*** | GFP-chsB::AFpyrG slab::thiAp-slab::AFriboB nkuAΔ::argB pyroA4 riboB2 pyrG89 | Martzoukou et al., 2018 |
| ***wt pall-gfp*** | pGEM-alcAp-Pall-GFP::panto pyrG89 pabaA1 argB nkuA? | Dimou et al., 2022 |
| ***ΔvapA pall-gfp*** | ΔvapA/ΔAN4406::AFpyrG pGEM-alcAp-Pall-GFP::pantoB pyrG89 pabaA1 nkuA? | This study |
| ***wt uapA-gfp*** | *ΔuapA::uapA-gfp::AFriboB ΔuapC::AfpyrG ΔnkuA::argB pabaA1 pyroA4 riboB2* | Evangelinos et al., 2016 |
| ***ΔvapA uapA-gfp*** | (AN4406)vapAΔ::AFpyroA uapAΔ::uapA-GFPL nkuAΔ::argB pyrG89 pyroA4 | This study |
| ***wt pmaA-gfp*** | TNO2A7 AN4859-(5xGA)GFP::AFpyrG nkuAΔ::argB pyrG89 riboB2 pyroA4 | Dimou et al., 2022 |
| ***ΔvapA pmaA-gfp*** | (AN4406)vapAΔ::AFpyrG AN4859-(5xGA)GFP::AFpyrG pantoB100 | This study |
| ***alcAp-gfp-oshB*** | oshB(p)-gfp-oshB(AN3424) pyroA4 | Buhler et al. |
| ***ΔvapA alcAp-gfp-oshB*** | (AN4406)vapAΔ::AFpyroA oshB(p)-gfp-oshB(AN3424) | This study |
| ***wt sac1-GFP*** | (AN3841)sac1-GFP::AFpyrG nkuAΔ::argB pyroA4 riboB2 pyrG89 | This study |
| ***ΔvapA sac1-GFP*** | (AN4406)vapAΔ::AFpyroA (AN3841)sac1-GFP::AFpyrG nkuAΔ::argB pyroA4 riboB2 pyrG89 | This study |
| ***thiAp-sac1 DnfA-GFP*** | DnfA-GFP::AFpyrG thiAp-AN3841::AFriboB nkuAΔ::argB pyrG89 pyroA4 riboB2 | This study |
| ***ΔvapA thiAp-sac1*** | (AN4406)vapAΔ::AFpyrG DnfA-GFP::AFpyrG thiAp-AN3841::AFriboB pantoB100 | This study |
| ***mCherry-vapA sac1-gfp*** | gpdAp-mCherry-(GA)x2-AN4406/vapA-trpC-panB (AN3841)sac1-GFP::AFpyrG pyrG89 argB2 pabaA1 pantoB100 | This study |

**Supplementary Table S2:** Annotations.

| **FungiDB ID** | **Systematic/Protein name** | **Cellular process** |
| --- | --- | --- |
| AN4406 | VapA/ Scs2/22 | ER-PM tethering |
| AN9149 | TcbA | ER-PM tethering |
| AN5624 | TcbB | ER-PM tethering |
| AN7165 | Ist2B | ER-PM tethering |
| AN2477 | Ist2A | ER-PM tethering |
| AN0722 | Ap2σ | Endocytosis |
| AN2756 | SlaB | Endocytosis |
| AN1023 | SagA | Endocytosis |
| AN12237 | AbpA | Endocytosis |
| AN1558 | MyoA | Endocytosis |
| AN7682 | Ap1σ/Aps1 | Late Golgi sorting |
| AN9526 | SedV/Sed5 | Qa-SNARE |
| AN8769 | SynA/Snc1/2 | R-SNARE |
| AN8672 | DnfA | apical cargo |
| AN6112 | DnfB | apical cargo |
| AN2523 | ChsB/Chs3 | apical cargo |
| AN4859 | PmaA | non-polar cargo |
| AN4853 | Pall | non-polar cargo |
| AN6932 | UapA | non-polar cargo |
| AN3424 | OshB/Osh3 | Oxysterol-binding |
| AN3841 | Sac1 | PI catabolism |
| AN1901 | Erg11A | Ergosterol biosynthesis |
| AN8283 | Erg11B | Ergosterol biosynthesis |
| AN2684 | Erg4A | Ergosterol biosynthesis |
| AN10648 | Erg4B | Ergosterol biosynthesis |
| AN0913 | PisA | PI biosynthesis |
| AN0640 | BasA | Sphingolipids biosynthesis |

**Supplementary Table S3:**
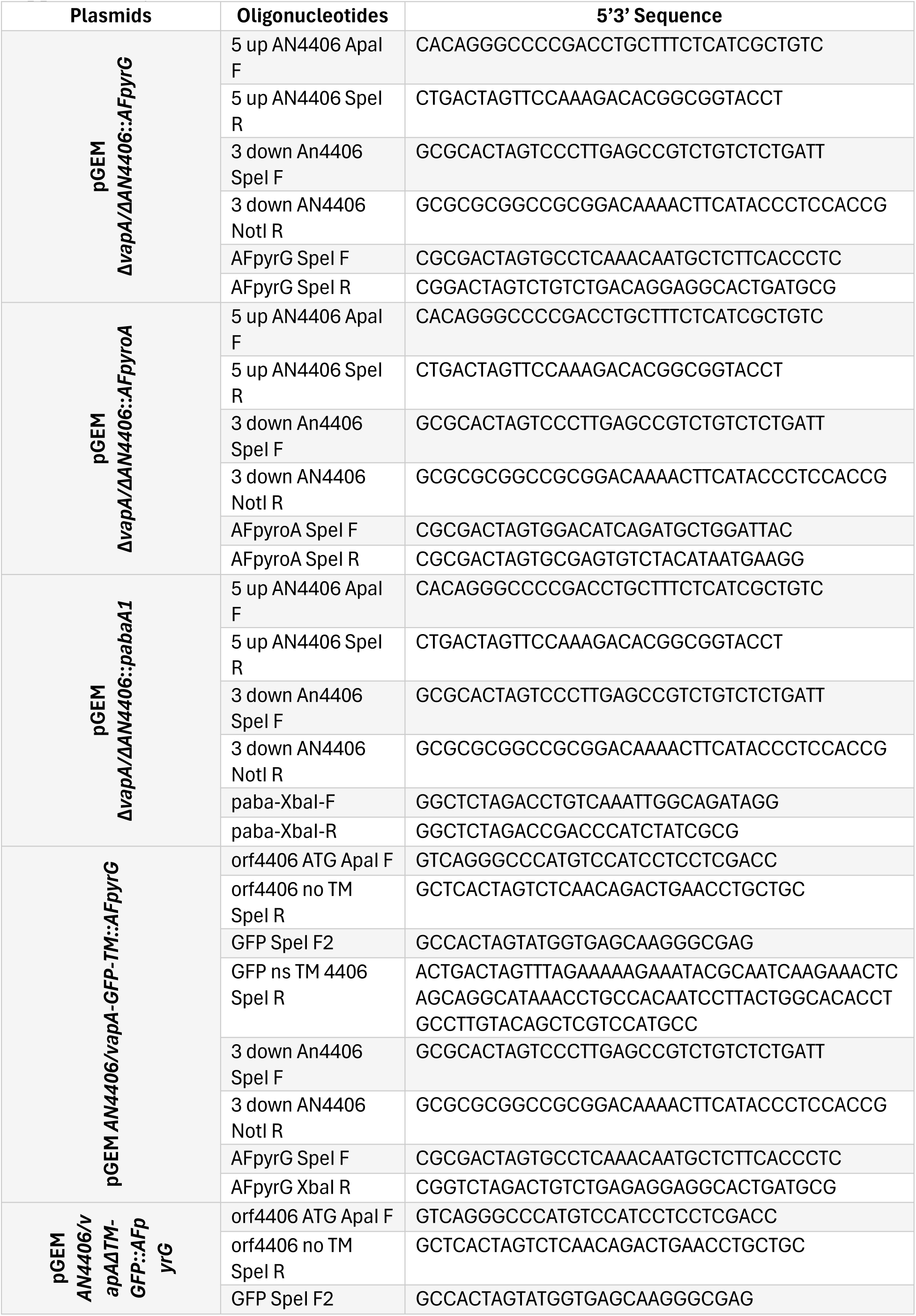
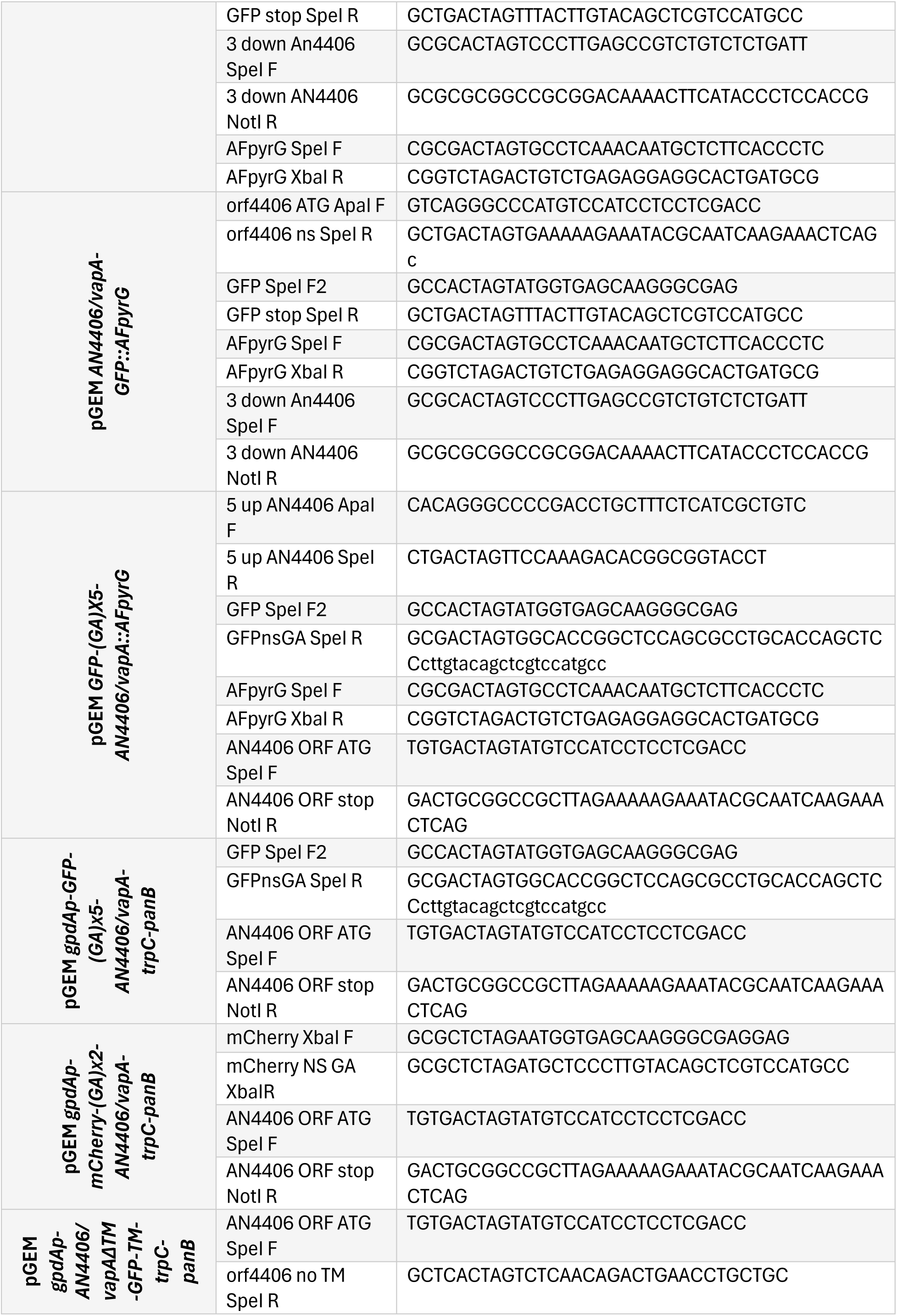
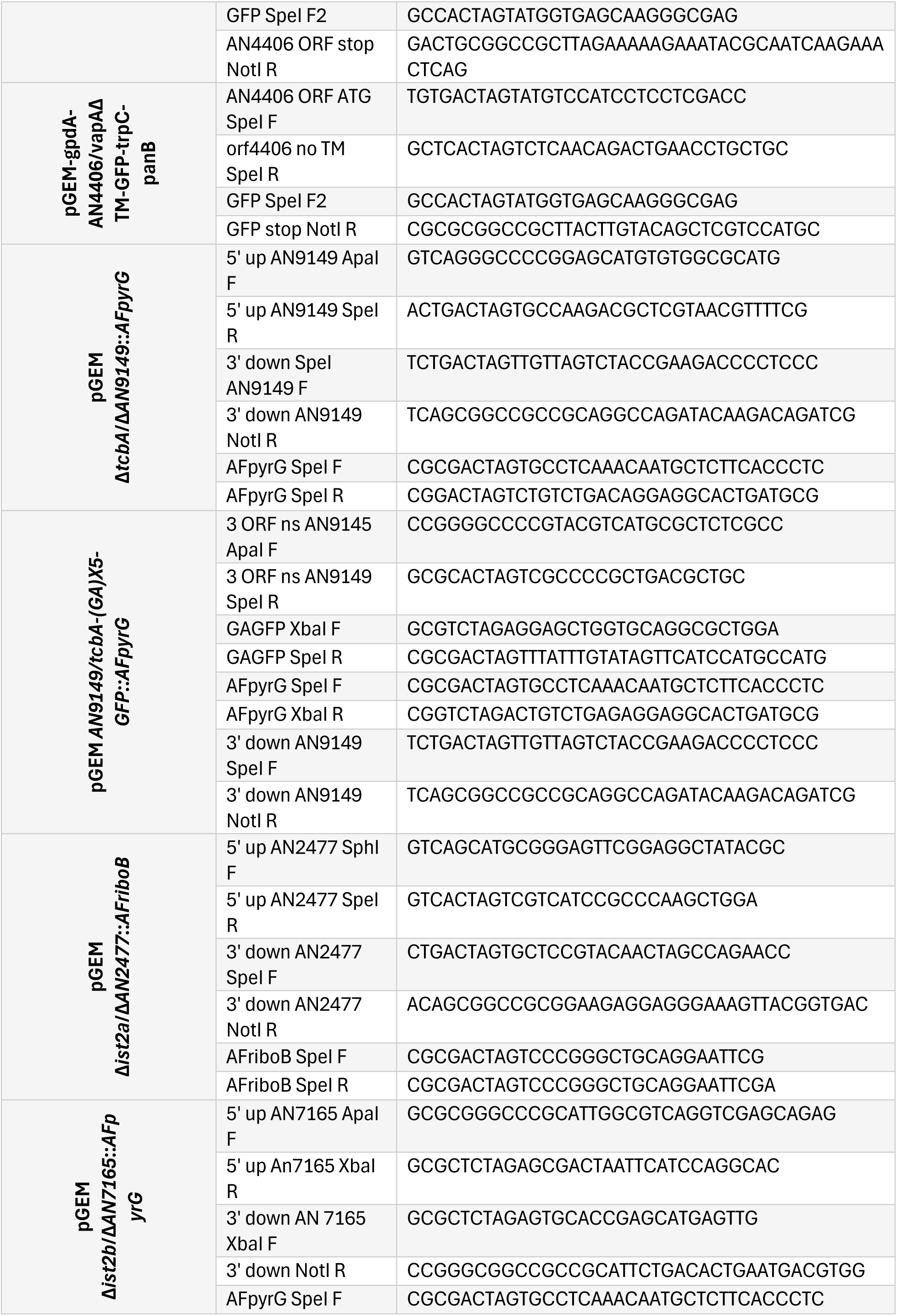
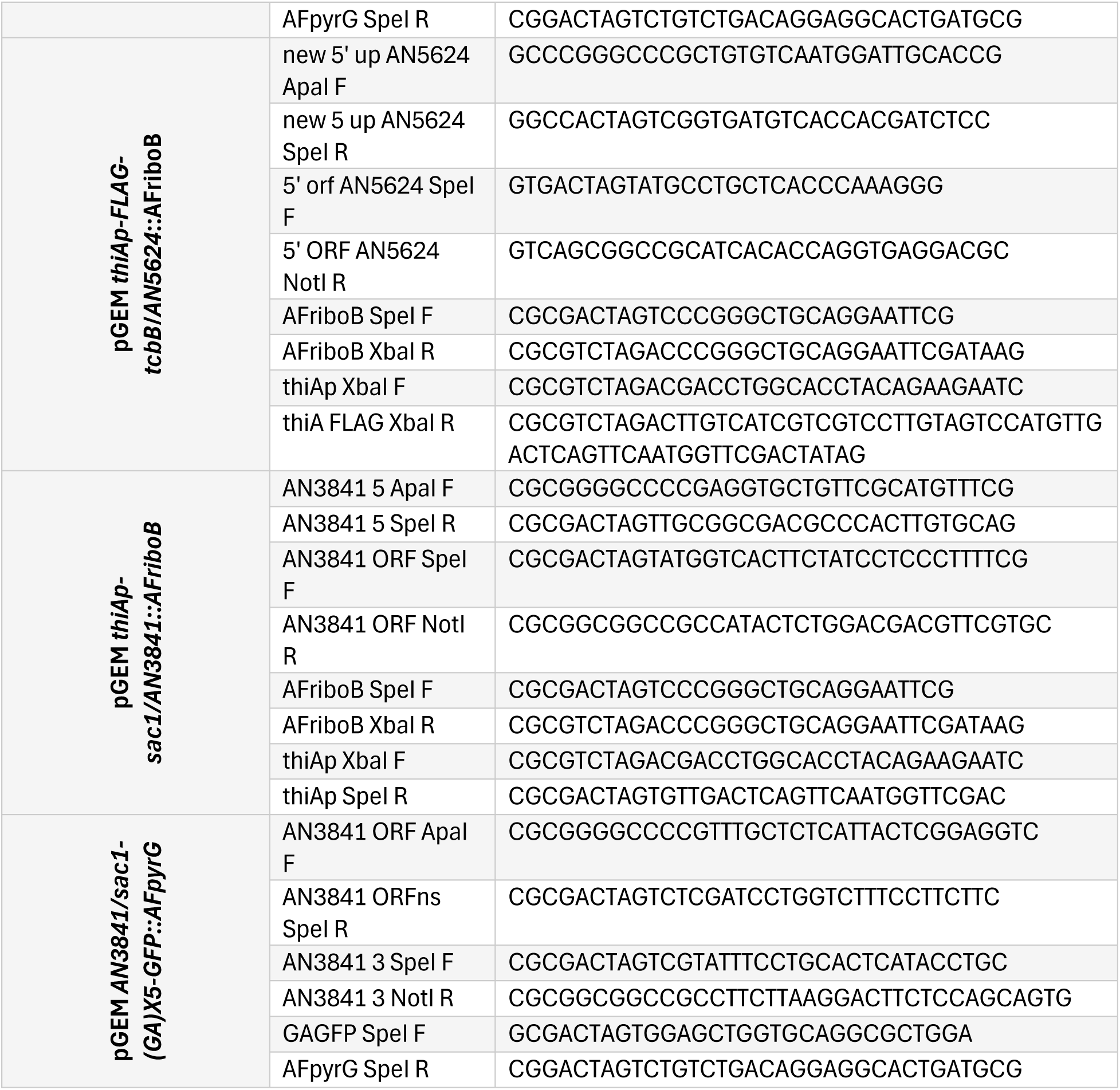
Primers.

**Supplementary Table S4:**
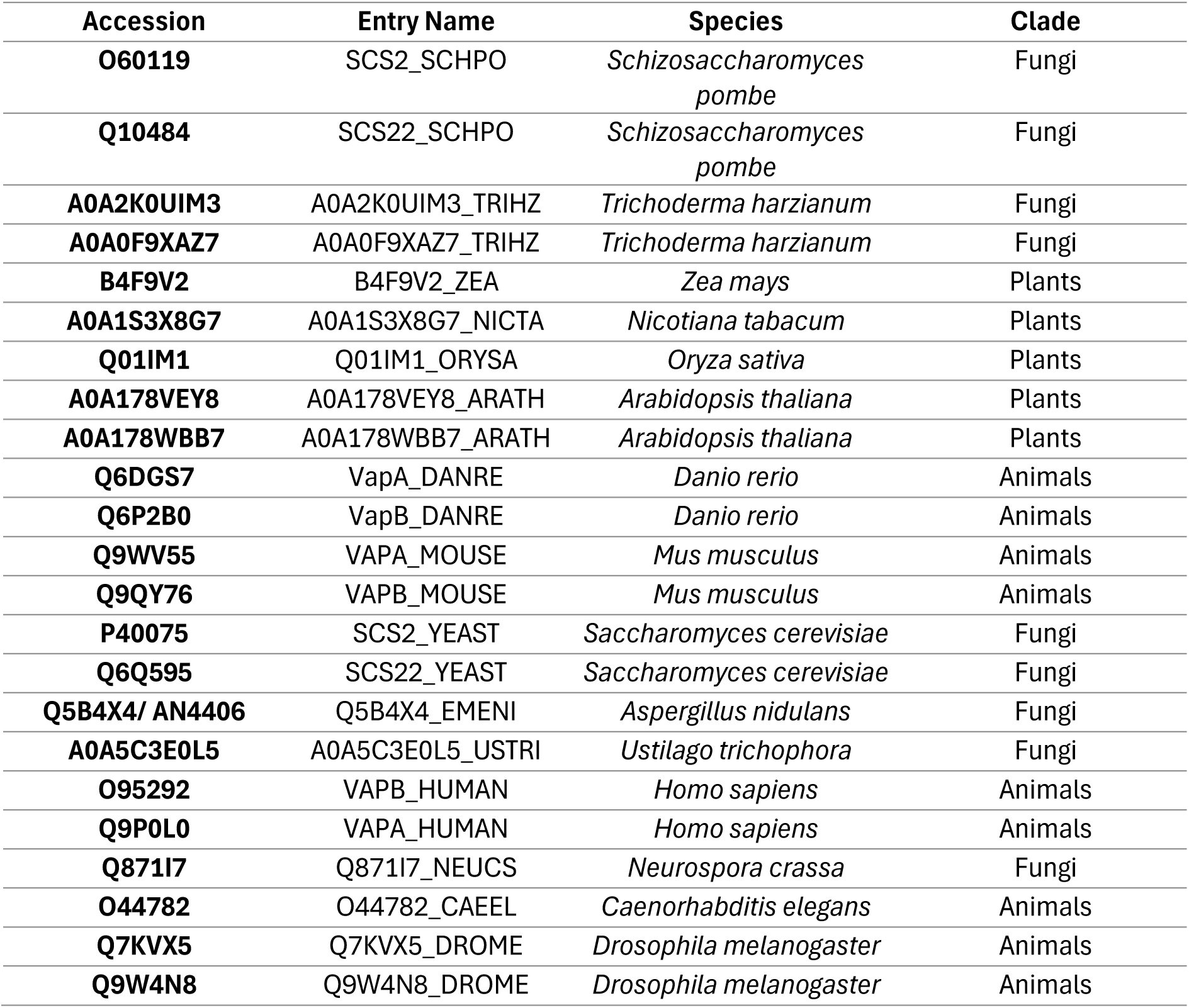
Phylogenetic tree IDs.

## REFERENCES

1. Phillips MJ, and Voeltz GK (2016). Structure and function of ER membrane contact sites with other organelles. Nat Rev Mol Cell Biol. 17(2): 69–82. doi: 10.1038/nrm.2015.8.

2. Pérez-Sancho J, Tilsner J, Samuels AL, Botella MA, Bayer EM, and Rosado A (2016). Stitching Organelles: Organization and Function of Specialized Membrane Contact Sites in Plants. Trends Cell Biol. 26(9): 705–717. doi: 10.1016/j.tcb.2016.05.007.

3. Scorrano L, De Matteis MA, Emr S, Giordano F, Hajnóczky G, Kornmann B, Lackner LL, Levine TP, Pellegrini L, Reinisch K, Rizzuto R, Simmen T, Stenmark H, Ungermann C, and Schuldiner M (2019). Coming together to define membrane contact sites. Nat Commun. 10(1): 1287. doi: 10.1038/s41467-019-09253-3.

4. Prinz WA, Toulmay A, and Balla T (2020). The functional universe of membrane contact sites. Nat Rev Mol Cell Biol. 21(1): 7–24. doi: 10.1038/s41580-019-0180-9.

5. Guillén-Samander A, and De Camilli P (2023). Endoplasmic Reticulum Membrane Contact Sites, Lipid Transport, and Neurodegeneration. Cold Spring Harb Perspect Biol. 15(4): a041257. doi: 10.1101/cshperspect.a041257.

6. Nishimura T, and Stefan CJ (2020). Specialized ER membrane domains for lipid metabolism and transport. Biochim Biophys Acta Mol Cell Biol Lipids. 1865(1): 158492. doi: 10.1016/j.bbalip.2019.07.001.

7. Jang W, and Haucke V (2024). ER remodeling via lipid metabolism. Trends Cell Biol. 34(11): 942–954. doi: 10.1016/j.tcb.2024.01.011.

8. Manford AG, Stefan CJ, Yuan HL, Macgurn JA, and Emr SD (2012). ER-to-plasma membrane tethering proteins regulate cell signaling and ER morphology. Dev Cell. 23(6): 1129–1140. doi: 10.1016/j.devcel.2012.11.004.

9. Henne WM, Liou J, and Emr SD (2015). Molecular mechanisms of inter-organelle ER–PM contact sites. Curr Opin Cell Biol. 35: 123–130. doi: 10.1016/j.ceb.2015.05.001.

10. Quon E, Sere YY, Chauhan N, Johansen J, Sullivan DP, Dittman JS, Rice WJ, Chan RB, Di Paolo G, Beh CT, and Menon AK (2018). Endoplasmic reticulum-plasma membrane contact sites integrate sterol and phospholipid regulation. PLoS Biol. 16(5): e2003864. doi: 10.1371/journal.pbio.2003864.

11. Dionysopoulou M, and Diallinas G (2021). Impact of Membrane Lipids on UapA and AzgA Transporter Subcellular Localization and Activity in Aspergillus nidulans. J Fungi Basel Switz. 7(7): 514. doi: 10.3390/jof7070514.

12. Fratti RA (2021). Editorial: Effects of Membrane Lipids on Protein Function. Front Cell Dev Biol. 9: 675264. doi: 10.3389/fcell.2021.675264.

13. Kors S, Costello JL, and Schrader M (2022). VAP Proteins - From Organelle Tethers to Pathogenic Host Interactors and Their Role in Neuronal Disease. Front Cell Dev Biol. 10: 895856. doi: 10.3389/fcell.2022.895856.

14. Liu D, Yuan H, Chen S, Ferro-Novick S, and Novick P (2024). Different ER-plasma membrane tethers play opposing roles in autophagy of the cortical ER. Proc Natl Acad Sci U S A. 121(24): e2321991121. doi: 10.1073/pnas.2321991121.

15. Whitlock JM, and Hartzell HC (2017). Anoctamins/TMEM16 Proteins: Chloride Channels Flirting with Lipids and Extracellular Vesicles. Annu Rev Physiol. 79(Volume 79, 2017): 119–143. doi: 10.1146/annurev-physiol-022516-034031.

16. Saheki Y, and De Camilli P (2017). The Extended-Synaptotagmins. Biochim Biophys Acta Mol Cell Res. 1864(9): 1490–1493. doi: 10.1016/j.bbamcr.2017.03.013.

17. Hoffmann PC, Bharat TAM, Wozny MR, Boulanger J, Miller EA, and Kukulski W (2019). Tricalbins Contribute to Cellular Lipid Flux and Form Curved ER-PM Contacts that Are Bridged by Rod-Shaped Structures. Dev Cell. 51(4): 488–502.e8. doi: 10.1016/j.devcel.2019.09.019.

18. Murphy SE, and Levine TP (2016). VAP, a Versatile Access Point for the Endoplasmic Reticulum: Review and analysis of FFAT-like motifs in the VAPome. Biochim Biophys Acta. 1861(8 Pt B): 952–961. doi: 10.1016/j.bbalip.2016.02.009.

19. Kodama TS, Furuita K, and Kojima C (2025). Beyond Static Tethering at Membrane Contact Sites: Structural Dynamics and Functional Implications of VAP Proteins. Mol Basel Switz. 30(6): 1220. doi: 10.3390/molecules30061220.

20. Collado J, Kalemanov M, Campelo F, Bourgoint C, Thomas F, Loewith R, Martínez-Sánchez A, Baumeister W, Stefan CJ, and Fernández-Busnadiego R (2019). Tricalbin-Mediated Contact Sites Control ER Curvature to Maintain Plasma Membrane Integrity. Dev Cell. 51(4): 476–487.e7. doi: 10.1016/j.devcel.2019.10.018.

21. Stefan CJ (2020). Endoplasmic reticulum-plasma membrane contacts: Principals of phosphoinositide and calcium signaling. Curr Opin Cell Biol. 63: 125–134. doi: 10.1016/j.ceb.2020.01.010.

22. Thomas FB, Omnus DJ, Bader JM, Chung GH, Kono N, and Stefan CJ (2022). Tricalbin proteins regulate plasma membrane phospholipid homeostasis. Life Sci Alliance. 5(8): e202201430. doi: 10.26508/lsa.202201430.

23. Wolf W, Kilic A, Schrul B, Lorenz H, Schwappach B, and Seedorf M (2012). Yeast Ist2 Recruits the Endoplasmic Reticulum to the Plasma Membrane and Creates a Ribosome-Free Membrane Microcompartment. PLOS ONE. 7(7): e39703. doi: 10.1371/journal.pone.0039703.

24. Giordano F, Saheki Y, Idevall-Hagren O, Colombo SF, Pirruccello M, Milosevic I, Gracheva EO, Bagriantsev SN, Borgese N, and De Camilli P (2013). PI(4,5)P2-Dependent and Ca2+-Regulated ER-PM Interactions Mediated by the Extended Synaptotagmins. Cell. 153(7): 1494–1509. doi: 10.1016/j.cell.2013.05.026.

25. Saheki Y, Bian X, Schauder CM, Sawaki Y, Surma MA, Klose C, Pincet F, Reinisch KM, and De Camilli P (2016). Control of plasma membrane lipid homeostasis by the extended synaptotagmins. Nat Cell Biol. 18(5): 504–515. doi: 10.1038/ncb3339.

26. Schauder CM, Wu X, Saheki Y, Narayanaswamy P, Torta F, Wenk MR, De Camilli P, and Reinisch KM (2014). Structure of a lipid-bound extended synaptotagmin indicates a role in lipid transfer. Nature. 510(7506): 552–555. doi: 10.1038/nature13269.

27. Yu H, Liu Y, Gulbranson DR, Paine A, Rathore SS, and Shen J (2016). Extended synaptotagmins are Ca2+-dependent lipid transfer proteins at membrane contact sites. Proc Natl Acad Sci U S A. 113(16): 4362–4367. doi: 10.1073/pnas.1517259113.

28. Creutz CE, Snyder SL, and Schulz TA (2004). Characterization of the yeast tricalbins: membrane-bound multi-C2-domain proteins that form complexes involved in membrane trafficking. Cell Mol Life Sci CMLS. 61(10): 1208–1220. doi: 10.1007/s00018-004-4029-8.

29. Schulz TA, and Creutz CE (2004). The tricalbin C2 domains: lipid-binding properties of a novel, synaptotagmin-like yeast protein family. Biochemistry. 43(13): 3987–3995. doi: 10.1021/bi036082w.

30. Toulmay A, and Prinz WA (2012). A conserved membrane-binding domain targets proteins to organelle contact sites. J Cell Sci. 125(Pt 1): 49–58. doi: 10.1242/jcs.085118.

31. Zhang D, Vjestica A, and Oliferenko S (2012). Plasma Membrane Tethering of the Cortical ER Necessitates Its Finely Reticulated Architecture. Curr Biol. 22(21): 2048–2052. doi: 10.1016/j.cub.2012.08.047.

32. Ng AYE, Ng AQE, and Zhang D (2018). ER-PM Contacts Restrict Exocytic Sites for Polarized Morphogenesis. Curr Biol CB. 28(1): 146–153.e5. doi: 10.1016/j.cub.2017.11.055.

33. Ng AQE, Ng AYE, and Zhang D (2020). Plasma Membrane Furrows Control Plasticity of ER-PM Contacts. Cell Rep. 30(5): 1434–1446.e7. doi: 10.1016/j.celrep.2019.12.098.

34. Hoh KL, Mu B, See T, Ng AYE, Ng AQE, and Zhang D (2024). VAP-mediated membrane-tethering mechanisms implicate ER-PM contact function in pH homeostasis. Cell Rep. 43(8). doi: 10.1016/j.celrep.2024.114592.

35. Zhang J, Chen X, Yang Z, Xu H, Weng S, Wang Z, and Tang W (2022). Endoplasmic reticulum membrane protein MoScs2 is important for asexual development and pathogenesis of Magnaporthe oryzae. Front Microbiol. 13: 906784. doi: 10.3389/fmicb.2022.906784.

36. Martzoukou O, Amillis S, Zervakou A, Christoforidis S, and Diallinas G (2017). The AP-2 complex has a specialized clathrin-independent role in apical endocytosis and polar growth in fungi. eLife. 6: e20083. doi: 10.7554/eLife.20083.

37. Martzoukou O, Diallinas G, and Amillis S (2018). Secretory Vesicle Polar Sorting, Endosome Recycling and Cytoskeleton Organization Require the AP-1 Complex in Aspergillus nidulans. Genetics. 209(4): 1121–1138. doi: 10.1534/genetics.118.301240.

38. Dimou S, Martzoukou O, Dionysopoulou M, Bouris V, Amillis S, and Diallinas G (2020). Translocation of nutrient transporters to cell membrane via Golgi bypass in Aspergillus nidulans. EMBO Rep. 21(7): e49929. doi: 10.15252/embr.201949929.

39. Markina-Iñarrairaegui A, Pantazopoulou A, Espeso EA, and Peñalva MA (2013). The Aspergillus nidulans Peripheral ER: Disorganization by ER Stress and Persistence during Mitosis. PLoS ONE. 8(6): e67154. doi: 10.1371/journal.pone.0067154.

40. Tojima T, Suda Y, Jin N, Kurokawa K, and Nakano A (2024). Spatiotemporal dissection of the Golgi apparatus and the ER-Golgi intermediate compartment in budding yeast. eLife.13. doi: 10.7554/elife.92900.

41. Pinar M, Pantazopoulou A, Arst HN, and Peñalva MA (2013). Acute inactivation of the *Aspergillus nidulans* Golgi membrane fusion machinery: correlation of apical extension arrest and tip swelling with cisternal disorganization. Mol Microbiol. 89(2): 228–248. doi: 10.1111/mmi.12280.

42. Pantazopoulou A, and Peñalva MA (2009). Organization and Dynamics of the*Aspergillus nidulans*Golgi during Apical Extension and Mitosis. Mol Biol Cell. 20(20): 4335–4347. doi: 10.1091/mbc.e09-03-0254.

43. Levine TP, and Munro S (2002). Targeting of Golgi-Specific Pleckstrin Homology Domains Involves Both PtdIns 4-Kinase-Dependent and -Independent Components. Curr Biol. 12(9): 695–704. doi: 10.1016/S0960-9822(02)00779-0.

44. Evangelinos M, Martzoukou O, Chorozian K, Amillis S, and Diallinas G (2016). BsdABsd2-dependent vacuolar turnover of a misfolded version of the UapA transporter along the secretory pathway: prominent role of selective autophagy. Mol Microbiol. 100(5): 893– 911. doi: 10.1111/mmi.13358.

45. Peñalva MA (2005). Tracing the endocytic pathway of Aspergillus nidulans with FM4-64. Fungal Genet Biol. 42(12): 963–975. doi: 10.1016/j.fgb.2005.09.004.

46. Stefan CJ, Manford AG, Baird D, Yamada-Hanff J, Mao Y, and Emr SD (2011). Osh Proteins Regulate Phosphoinositide Metabolism at ER-Plasma Membrane Contact Sites. Cell. 144(3): 389–401. doi: 10.1016/j.cell.2010.12.034.

47. Apostolaki A, Harispe L, Calcagno-Pizarelli AM, Vangelatos I, Sophianopoulou V, Arst Jr HN, Peñalva MA, Amillis S, and Scazzocchio C (2012). *Aspergillus nidulans* CkiA is an essential casein kinase I required for delivery of amino acid transporters to the plasma membrane. Mol Microbiol. 84(3): 530–549. doi: 10.1111/j.1365-2958.2012.08042.x.

48. Roy A, and Levine TP (2004). Multiple Pools of Phosphatidylinositol 4-Phosphate Detected Using the Pleckstrin Homology Domain of Osh2p. J Biol Chem. 279(43): 44683– 44689. doi: 10.1074/jbc.m401583200.

49. Bühler N, Hagiwara D, and Takeshita N (2015). Functional Analysis of Sterol Transporter Orthologues in the Filamentous Fungus Aspergillus nidulans. Eukaryot Cell. 14(9): 908– 921. doi: 10.1128/ec.00027-15.

50. Ramón A, Muro-Pastor MI, Scazzocchio C, and Gonzalez R (2000). Deletion of the unique gene encoding a typical histone H1 has no apparent phenotype in *Aspergillus nidulans*. Mol Microbiol. 35(1): 223–233. doi: 10.1046/j.1365-2958.2000.01702.x.

51. Dimou S, Kourkoulou A, Amillis S, Percudani R, and Diallinas G (2019). The peroxisomal SspA protein is redundant for purine utilization but essential for peroxisome localization in septal pores in Aspergillus nidulans. Fungal Genet Biol. 132: 103259. doi: 10.1016/j.fgb.2019.103259.

52. Galanopoulou K, Scazzocchio C, Galinou ME, Liu W, Borbolis F, Karachaliou M, Oestreicher N, Hatzinikolaou DG, Diallinas G, and Amillis S (2014). Purine utilization proteins in the Eurotiales: cellular compartmentalization, phylogenetic conservation and divergence. Fungal Genet Biol FG B. 69: 96–108. doi: 10.1016/j.fgb.2014.06.005.

53. Lemmon MA (2004). Pleckstrin homology domains: not just for phosphoinositides. Biochem Soc Trans. 32(5): 707–711. doi: 10.1042/bst0320707.

54. Kourkoulou A, Martzoukou O, Fischer R, and Amillis S (2024). A type II phosphatidylinositol-4-kinase coordinates sorting of cargo polarizing by endocytic recycling. Commun Biol. 7(1): 855. doi: 10.1038/s42003-024-06553-3.

55. Schultzhaus Z, Zheng W, Wang Z, Mouriño-Pérez R, and Shaw B (2017). Phospholipid flippases DnfA and DnfB exhibit differential dynamics within the A. nidulans Spitzenkörper. Fungal Genet Biol. 99: 26–28. doi: 10.1016/j.fgb.2016.12.007.

56. Fukuda K, Yamada K, Deoka K, Yamashita S, Ohta A, and Horiuchi H (2009). Class III Chitin Synthase ChsB of *Aspergillus nidulans* Localizes at the Sites of Polarized Cell Wall Synthesis and Is Required for Conidial Development. Eukaryot Cell. 8(7): 945–956. doi: 10.1128/ec.00326-08.

57. López-Hernández T, Takenaka K, Mori Y, Kongpracha P, Nagamori S, Haucke V, and Takamori S (2022). Clathrin-independent endocytic retrieval of SV proteins mediated by the clathrin adaptor AP-2 at mammalian central synapses. eLife. 11. doi: 10.7554/elife.71198.

58. Alvarez FJ, Douglas LM, and Konopka JB (2007). Sterol-Rich Plasma Membrane Domains in Fungi. Eukaryot Cell. 6(5): 755–763. doi: 10.1128/ec.00008-07.

59. Takeshita N, Higashitsuji Y, Konzack S, and Fischer R (2008). Apical Sterol-rich Membranes Are Essential for Localizing Cell End Markers That Determine Growth Directionality in the Filamentous Fungus*Aspergillus nidulans*. Mol Biol Cell. 19(1): 339–351. doi: 10.1091/mbc.e07-06-0523.

60. Lu R, Drubin DG, and Sun Y (2016). Clathrin-mediated endocytosis in budding yeast at a glance. J Cell Sci. 129(8): 1531–1536. doi: 10.1242/jcs.182303.

61. Hill JM, Pedersen RT, and Drubin DG (2024). Myosin-I’s motor and actin assembly activation activities are modular and separable in budding yeast clathrin-mediated endocytosis. MicroPublication Biol. 2024. doi: 10.17912/micropub.biology.001223.

62. Araujo-Bazán L, Peñalva MA, and Espeso EA (2008). Preferential localization of the endocytic internalization machinery to hyphal tips underlies polarization of the actin cytoskeleton in *Aspergillus nidulans*. Mol Microbiol. 67(4): 891–905. doi: 10.1111/j.1365-2958.2007.06102.x.

63. Taheri-Talesh N, Horio T, Araujo-Bazán L, Dou X, Espeso EA, Peñalva MA, Osmani SA, and Oakley BR (2008). The Tip Growth Apparatus of *Aspergillus nidulans*. Mol Biol Cell. 19(4): 1439–1449. doi: 10.1091/mbc.e07-05-0464.

64. Taheri-Talesh N, Xiong Y, and Oakley BR (2012). The Functions of Myosin II and Myosin V Homologs in Tip Growth and Septation in Aspergillus nidulans. PLoS ONE. 7(2): e31218. doi: 10.1371/journal.pone.0031218.

65. Hervás-Aguilar A, and Peñalva MA (2010). Endocytic Machinery Protein SlaB Is Dispensable for Polarity Establishment but Necessary for Polarity Maintenance in Hyphal Tip Cells of Aspergillus nidulans. Eukaryot Cell. 9(10): 1504–1518. doi: 10.1128/EC.00119-10.

66. Hernández-González M, Bravo-Plaza I, Pinar M, Ríos V de los, Jr HNA, and Peñalva MA (2018). Endocytic recycling via the TGN underlies the polarized hyphal mode of life. PLOS Genet. 14(4): e1007291. doi: 10.1371/journal.pgen.1007291.

67. Steinberg G, Peñalva MA, Riquelme M, Wösten HA, and Harris SD (2017). Cell Biology of Hyphal Growth. Microbiol Spectr. 5(2). doi: 10.1128/microbiolspec.funk-0034-2016.

68. Dimou S, Dionysopoulou M, Sagia GM, and Diallinas G (2022). Golgi-Bypass Is a Major Unconventional Route for Translocation to the Plasma Membrane of Non-Apical Membrane Cargoes in Aspergillus nidulans. Front Cell Dev Biol. 10. doi: 10.3389/fcell.2022.852028.

69. Sagia GM, Georgiou X, Chamilos G, Diallinas G, and Dimou S (2024). Distinct trafficking routes of polarized and non-polarized membrane cargoes in Aspergillus nidulans. eLife. 13: e103355. doi: 10.7554/eLife.103355.

70. Georgiou X, Dimou S, Diallinas G, and Samiotaki M (2023). The interactome of the UapA transporter reveals putative new players in anterograde membrane cargo trafficking. Fungal Genet Biol FG B. 169: 103840. doi: 10.1016/j.fgb.2023.103840.

71. Pantazopoulou A, and Diallinas G (2007). Fungal nucleobase transporters. FEMS Microbiol Rev. 31(6): 657–675. doi: 10.1111/j.1574-6976.2007.00083.x.

72. Kourkoulou A, Zantza I, Foti K, Mikros E, and Diallinas G (2021). Context-dependent Cryptic Roles of Specific Residues in Substrate Selectivity of the UapA Purine Transporter. J Mol Biol. 433(16): 166814. doi: 10.1016/j.jmb.2021.166814.

73. Krypotou E, and Diallinas G (2014). Transport assays in filamentous fungi: Kinetic characterization of the UapC purine transporter of *Aspergillus nidulans*. Fungal Genet Biol. 63: 1–8. doi: 10.1016/j.fgb.2013.12.004.

74. Gournas C, Amillis S, Vlanti A, and Diallinas G (2010). Transport-dependent endocytosis and turnover of a uric acid-xanthine permease. Mol Microbiol. 75(1): 246–260. doi: 10.1111/j.1365-2958.2009.06997.x.

75. Nishimura AL, Mitne-Neto M, Silva HCA, Richieri-Costa A, Middleton S, Cascio D, Kok F, Oliveira JRM, Gillingwater T, Webb J, Skehel P, and Zatz M (2004). A Mutation in the Vesicle-Trafficking Protein VAPB Causes Late-Onset Spinal Muscular Atrophy and Amyotrophic Lateral Sclerosis. Am J Hum Genet. 75(5): 822–831. doi: 10.1086/425287.

76. De Vos KJ, Mórotz GM, Stoica R, Tudor EL, Lau K-F, Ackerley S, Warley A, Shaw CE, and Miller CCJ (2012). VAPB interacts with the mitochondrial protein PTPIP51 to regulate calcium homeostasis. Hum Mol Genet. 21(6): 1299–1311. doi: 10.1093/hmg/ddr559.

77. Paillusson S, Gomez-Suaga P, Stoica R, Little D, Gissen P, Devine MJ, Noble W, Hanger DP, and Miller CCJ (2017). α-Synuclein binds to the ER–mitochondria tethering protein VAPB to disrupt Ca2+ homeostasis and mitochondrial ATP production. Acta Neuropathol (Berl). 134(1): 129–149. doi: 10.1007/s00401-017-1704-z.

78. Dudás EF, Huynen MA, Lesk AM, and Pastore A (2021). Invisible leashes: The tethering VAPs from infectious diseases to neurodegeneration. J Biol Chem. 296: 100421. doi: 10.1016/j.jbc.2021.100421.

79. Stefano G, Renna L, Wormsbaecher C, Gamble J, Zienkiewicz K, and Brandizzi F (2018). Plant Endocytosis Requires the ER Membrane-Anchored Proteins VAP27-1 and VAP27-3. Cell Rep. 23(8): 2299–2307. doi: 10.1016/j.celrep.2018.04.091.

80. Wang P, Duckney P, Gao E, Hussey PJ, Kriechbaumer V, Li C, Zang J, and Zhang T (2023). Keep in contact: multiple roles of endoplasmic reticulum–membrane contact sites and the organelle interaction network in plants. New Phytol. 238(2): 482–499. doi: 10.1111/nph.18745.

81. Riquelme M, Aguirre J, Bartnicki-García S, Braus GH, Feldbrügge M, Fleig U, Hansberg W, Herrera-Estrella A, Kämper J, Kück U, Mouriño-Pérez RR, Takeshita N, and Fischer R (2018). Fungal Morphogenesis, from the Polarized Growth of Hyphae to Complex Reproduction and Infection Structures. Microbiol Mol Biol Rev. 82(2). doi: 10.1128/mmbr.00068-17.

82. Barata-Antunes C, Talaia G, Broutzakis G, Ribas D, De Beule P, Casal M, Stefan CJ, Diallinas G, and Paiva S (2022). Interactions of cytosolic tails in the Jen1 carboxylate transporter are critical for trafficking and transport activity. J Cell Sci. 135(10). doi: 10.1242/jcs.260059.

83. Karachaliou M, Amillis S, Evangelinos M, Kokotos AC, Yalelis V, and Diallinas G (2013). The arrestin-like protein ARTA is essential for ubiquitination and endocytosis of the UAPA transporter in response to both broad-range and specific signals. Mol Microbiol. 88(2): 301–317. doi: 10.1111/mmi.12184.

84. Sioupouli G, Lambrinidis G, Mikros E, Amillis S, and Diallinas G (2017). Cryptic purine transporters in *Aspergillus nidulans* reveal the role of specific residues in the evolution of specificity in the NCS1 family. Mol Microbiol. 103(2): 319–332. doi: 10.1111/mmi.13559.

85. Nayak T, Szewczyk E, Oakley CE, Osmani A, Ukil L, Murray SL, Hynes MJ, Osmani SA, and Oakley BR (2006). A versatile and efficient gene-targeting system for Aspergillus nidulans. Genetics. 172(3): 1557–1566. doi: 10.1534/genetics.105.052563.

86. Krypotou E, Evangelidis T, Bobonis J, Pittis AA, Gabaldón T, Scazzocchio C, Mikros E, and Diallinas G (2015). Origin, diversification and substrate specificity in the family of NCS1/FUR transporters. Mol Microbiol. 96(5): 927–950. doi: 10.1111/mmi.12982.

87. Katoh K, and Standley DM (2013). MAFFT Multiple Sequence Alignment Software Version 7: Improvements in Performance and Usability. Mol Biol Evol. 30(4): 772–780. doi: 10.1093/molbev/mst010.

88. Nguyen L-T, Schmidt HA, Von Haeseler A, and Minh BQ (2015). IQ-TREE: A Fast and Effective Stochastic Algorithm for Estimating Maximum-Likelihood Phylogenies. Mol Biol Evol. 32(1): 268–274. doi: 10.1093/molbev/msu300.

89. Trifinopoulos J, Nguyen L-T, von Haeseler A, and Minh BQ (2016). W-IQ-TREE: a fast online phylogenetic tool for maximum likelihood analysis. Nucleic Acids Res. 44(W1): W232–W235. doi: 10.1093/nar/gkw256.

90. Kalyaanamoorthy S, Minh BQ, Wong TKF, Von Haeseler A, and Jermiin LS (2017). ModelFinder: fast model selection for accurate phylogenetic estimates. Nat Methods. 14(6): 587–589. doi: 10.1038/nmeth.4285.

91. Hoang DT, Chernomor O, Von Haeseler A, Minh BQ, and Vinh LS (2018). UFBoot2: Improving the Ultrafast Bootstrap Approximation. Mol Biol Evol. 35(2): 518–522. doi: 10.1093/molbev/msx281.

92. Schindelin J, Arganda-Carreras I, Frise E, Kaynig V, Longair M, Pietzsch T, Preibisch S, Rueden C, Saalfeld S, Schmid B, Tinevez J-Y, White DJ, Hartenstein V, Eliceiri K, Tomancak P, and Cardona A (2012). Fiji: an open-source platform for biological-image analysis. Nat Methods. 9(7): 676–682. doi: 10.1038/nmeth.2019.

93. McKinney W (2010). Data Structures for Statistical Computing in Python. In: Proc. Python Sci. Conf. SciPy, Austin, Texas; pp 56–61.

94. Virtanen P et al. (2020). SciPy 1.0: fundamental algorithms for scientific computing in Python. Nat Methods. 17(3): 261–272. doi: 10.1038/s41592-019-0686-2.

95. Harris CR, Millman KJ, Van Der Walt SJ, Gommers R, Virtanen P, Cournapeau D, Wieser E, Taylor J, Berg S, Smith NJ, Kern R, Picus M, Hoyer S, Van Kerkwijk MH, Brett M, Haldane A, Del Río JF, Wiebe M, Peterson P, Gérard-Marchant P, Sheppard K, Reddy T, Weckesser W, Abbasi H, Gohlke C, and Oliphant TE (2020). Array programming with NumPy. Nature. 585(7825): 357–362. doi: 10.1038/s41586-020-2649-2.

96. Bolte S, and Cordelières FP (2006). A guided tour into subcellular colocalization analysis in light microscopy. J Microsc. 224(3): 213–232. doi: 10.1111/j.1365-2818.2006.01706.x.

